# Hyperactive STAT1 Promotes T Follicular Helper Type 1 Cell Differentiation to Trigger Autoimmunity

**DOI:** 10.64898/2026.03.19.713058

**Authors:** Ran Chen, Xuemei Chen, Jigui Yang, Huilin Mu, Shengqiao Mao, Siyu Chen, Rui Gan, Qinglv Wei, Wenjing Tang, Junfeng Wu, Wuyang He, Satoshi Okada, Lina Zhou, Yunfei An, Xiaodong Zhao, Yanjun Jia

## Abstract

Heterozygous gain-of-function (GOF) mutations in signal transducer and activator of transcription 1 (*STAT1*) cause an inborn error of immunity characterized by immune dysregulation, recurrent infections and various autoimmune manifestations. However, the precise pathogenic mechanism by which *STAT1* GOF contributes to autoimmunity remains elusive. In our cohort, *STAT1*-GOF patients exhibit biased circulating follicular helper T (cTfh) populations with CXCR3^+^ Tfh1-like features. Using a *Stat1* GOF mouse model that spontaneously developed autoimmunity, we found that overactivated STAT1 promotes Tfh differentiation and disrupted T cell-dependent humoral responses with skewed immunoglobulin class switching towards IgG2. Furthermore, STAT1 GOF directly targets to Tfh and Th1 cell signature genes and thereby drives the development of Tfh1 cells with excessive IFN-γ production, which implicated in autoantibody production and the development of autoimmunity. Notably, IFN-γ neutralization significantly alleviated autoimmune cellular responses and autoantibody levels in mutant mice, highlighting IFN-γ blockade as a promising targeted therapy for the *STAT1*-GOF patients with autoimmunity. Our findings suggest that proper regulation of STAT1 activity within a reasonable magnitude is crucial for ensuring optimal host-protective humoral immunity.

**One-sentence summary:** Overactivated STAT1 promotes Tfh1 differentiation to drive autoimmunity.

## INTRODUCTION

Follicular helper T (Tfh) cells, originally characterized by high expression of the CXC-chemokine receptor 5 (CXCR5)(*1–4*), are a subset of effector CD4^+^T cells that is specialized in promoting the T cell-dependent B cell response and are essential for the formation and maintenance of germinal center (GC)(*5–7*). The canonical Tfh cell differentiation process initials at dendritic cell priming of a naïve CD4^+^ T cell in the T cell zone(*8–11*). Antigen stimulation upregulates the expression of transcription factor Bcl-6(*12–14*), inducible T cell co-stimulator (ICOS)(*9*), programmed cell death protein 1 (PD-1) (*15*) and CXCR5 while downregulating C-C chemokine receptor type 7(CCR7)and P-selectin glycoprotein ligand 1 (PSGL1), which coordinately guide activated CD4^+^ T cells to migrate toward the border of B cell follicles (T-B border) and away from the T cell zone. Upon recognizing cognate antigen-presenting B cells, the differentiating Tfh cells with strengthened Bcl-6 expression migrate deep into the interior of GC and further differentiate into fully functional Tfh cells, which assist in the production of high affinity class-switched antibodies, long-lived plasma cells and memory B cells(*5–7, 11*).

In addition to being critical for optimal protective immunity in response to infection and vaccination, Tfh cells have been linked to autoimmunity in both humans and animals(*7, 16–20*). Emerging evidence suggests that the abundance of circulating Tfh (cTfh) cells is increased in patients with various autoimmune diseases, including systemic lupus erythematosus (SLE)(*21, 22*), juvenile dermatomyositis (*23*), rheumatoid arthritis(*21*), type 1 diabetes(*24*), myasthenia gravis(*25*), Sjögren′s syndrome(*26*), systemic sclerosis(*27*), autoimmune hepatitis and autoimmune thyroid disease(*19, 28, 29*). Moreover, the elevated level of cTfh cells or specifical cTfh subgroup is strongly associated with the disease activity score. In mice, dysregulated accumulation of Tfh cells is found in lupus-prone *Roquin^san/san^* mice(*30, 31*), SLE-like BXSB-Yaa mice(*18*), autoimmune hepatitis-prone *Pdcd1*^-/-^mice(*19*), and autoimmune -prone Sle1 and NZB/W F1 mice(*32*). Furthermore, the dysregulated differentiation of Tfh cells may lead to the development of autoimmune diseases and are necessary for pathogenic autoantibody production(*16, 29, 33, 34*). Despite the potential significance, much remains unknown about the mechanics that contribute to Tfh cell dysregulation.

Signal transducer and activator of transcription (STAT)-mediated cytokine signaling pathways have been implicated in the Tfh cells development. STAT3-deficient CD4^+^ T cells show profound defects in Tfh cell differentiation, accompanied by decreased GC B cells and antigen-specific antibody production in response to immune challenge(*35, 36*). Loss-of function mutation in *STAT3* abolish IL-6/IL-21/IL-12-induced IL-21 expression in human CD4^+^ T cells, which compromised Tfh cell generation and abolished B-cell helper activity(*37*). In contrast, mice with STAT5 deficiency in CD4^+^ T cells exhibit an increase in Tfh and GC B cells and impairment of B-cell tolerance(*38, 39*). In further, previous work has shown that Tfh cells require STAT4 for IL-21 production but not for formation upon virus infection(*40*). STAT1 is a latent cytoplasmic transcription factor that mediates cell signaling in response to type I, II interferon (IFNs) and, to a lesser extent, IL-6. Early *in vitro* study demonstrated that type I IFN (IFN-α/β) enables STAT1 binding in the *Bcl6* locus to contribute to Tfh development(*41*). Yet, using adoptive transfers of TCR transgenic CD4^+^ T cells, others *in vivo* reported that STAT1 was merely required for early Tfh development, and there were no differences in Tfh cells or Bcl-6 expression at day 8 following virus challenging(*36, 42*). However, with STAT1-S727A mutant mice which the phosphorylation site of serine at position 727 is replaced by alanine, another study manifested that STAT1-pS727 activity is not necessary for Tfh and GC responses to foreign pathogen but can significantly alleviate autoimmunity as well as the associated Tfh cell expansion(*43*). Thus, the role of STAT1 in Tfh cell differentiation has been contentious, and needed to be further elucidated.

Heterozygous gain-of-function (GOF) mutations in *STAT1* locus were initially identified in patients with chronic mucocutaneous candidiasis (CMC) resulting from impaired Th17 differentiation(*44, 45*). Other than CMC, STAT1-GOF patients are susceptible to various infectious diseases and develop autoimmune manifestations(*46*). Several works revealed that cTfh in *STAT1*-GOF patients were skewed toward Th1-like cells despite similar frequency compared with health control(*37, 47*). Nevertheless, whether and how overactivated STAT1 regulates the differentiation of Tfh cells remains unknown. Although two group recently reported that aged mice with *Stat1* mutant (R274Q, T385M) exhibit autoimmune features(*47, 48*), the precise pathogenic mechanism by which STAT1 GOF contributes to disease is also poorly understood.

In this study, we generated a mouse model of STAT1 GOF (*Stat1*^T385M/+^ mice) that recapitulates human STAT1 GOF to dissect its role in Tfh biology and autoimmunity. We found that *Stat1*^T385M/+^ mice exhibited multiorgan inflammation and spontaneous autoimmune phenotype, with excessive Tfh accumulation and spontaneous GC formation. Nonetheless, *Stat1*^T385M/+^ mice displayed a defective T cell-dependent humoral immunity, and impaired in class-switched antigen-specific antibody responses. Further, hyperactive STAT1 intrinsically promoted the differentiation of IFN γ-producing Tfh cells with Th1 features, which induced the production of autoantibody and the development of autoimmunity. Mechanistically, genome-wide analysis indicated that *Stat1* T385M intensified its transcriptionally regulatory activity but did not change the profile of STAT1 targets.

Overactivated STAT1 directly targets both Tfh- and Th1-relevant signature genes to constantly promote the differentiation of Tfh1 cells. Importantly, similar to inhibit JAKs activity *in vivo,* IFN-γ neutralization can almost completely alleviate autoimmunity in *Stat1^Mut/WT^* mice. The findings imply that IFN-γ neutralization may be a novel treatment options for patients with *STAT1* GOF and other related autoimmune diseases.

## Materials and Methods

### Study design

The goal of this study is to elucidate how *STAT1* GOF mutation contribute to autoimmunity. With a cohort of *STAT1*-GOF patients from multiple regions of China, we analyzed the circulating follicular helper T cells in the peripheral blood and evaluated the autoimmune manifestations. In addition, we generated a corresponding STAT1 GOF mutant mouse model (*Stat1* T385M) and assessed the development of autoimmunity and the antibody levels. Further, we analyzed the effects of STAT1 GOF on the Tfh differentiation and the T cell-dependent humoral responses in various immunogens challenge. With adoptive transfer, bone marrow chimera and in vitro co-culture, we explored whether and how STAT1 GOF led to autoimmunity by promoting the differentiation of pathogenic Tfh cells. with RNA-seq and cleavage under targets and tagmentation (CUT&Tag) coupled with high-throughput sequencing, we further investigated how STAT1 GOF promoted the development of pathogenic Tfh cells. Based on the aforementioned study, we determined whether targeting pathogenic Tfh differentiation can alleviate or reverse the autoimmunity caused by STAT1 GOF *in vitro* and *in vivo*. Both male and female mice were used for experiments. All experimental analyses were nonblinded. Numbers of both experimental replicates and independent experiments were described in detail in the figure legends.

### Human

20 *STAT1*-GOF patients and 17 age-matched health controls were enrolled in this study. Human peripheral blood mononuclear cells (PBMCs) were isolated using Ficoll-Hypaque (GE Healthcare, USA) gradient centrifugation as previously described(*91*). The absolute number of total lymphocytes was determined by an automatic hematology analyzer (Sysmex XE-2100, Sysmex Co., Ltd, Japan). Written informed consent was obtained from all the subjects included in this study and their legal guardians in accordance with the Declaration of Helsinki and with approval from the Institutional Review Board of Children’s Hospital of Chongqing Medical University (2021–138).

### Mice

C57BL/6(B6), B6.SJL-PtprcaPepcb/BoyJ(B6.CD45.1) and B6.Cg-Tg(TcraTcrb)425Cbn/J (OTII) were originally purchased from The Jackson Laboratories.

C57BL/6JGpt-Rag1em1Cd3259/Gpt(Rag1 KO) was obtained from GemPharmatech Co., Ltd (China). Mutant STAT1 (p.T385M) mice carrying a missense mutation (c.1154C>T) were generated by in situ gene targeting using TurboKnockout^®^ technology (Cyagen Biosciences, China). The targeting vector was constructed, electroporated into ES cells, and positive clones were identified by PCR and Southern blot. Verified ES cells were injected into blastocysts, and chimeric mice were bred to obtain germline-transmitted mutants. Genomic DNA was extracted from tail biopsies for genotyping. A 539-bp fragment was amplified using the following primers: forward, 5′- TGTCTTCTGTGTGTTGAAGCGTAA-3′; reverse, 5′- ATAAAGCACTCTACTGAGGTCCG-3′. PCR products were analyzed by Sanger sequencing (Sangon, Shanghai) to confirm the mutation. Mice were categorized into three genotypes: wild-type (WT), heterozygous (*Stat1^Mut/WT^*), and homozygous (*Stat1^Mut/Mut^*). Littermates of different genotypes were used for comparative analyses in all experiments.

All mice used in this study were on a C57BL/6 background and were housed under specific pathogen-free conditions. All animal experiments were performed with 6- to 8-week-old mice, unless other indicated. Experimental analyses were nonblinded. All experimental procedures involving animals approved by the Laboratory Animal Welfare and Ethics Committee of the Children’s Hospital of Chongqing Medical University.

### Sanger sequencing

cDNA of P6 and P7 were amplified using primers as follow: P6: forward, 5′-TGTCTTCTGTGTGTTGAAGCGTAA-3′; reverse, 5′-GAAGGTGCGGTCCCATAACA-3′. P7: forward, 5′-TGAAGACAGGGGTCCAGTTC-3′; reverse, 5′-TGGCGTTAGGACCAAGAAGC-3′).

The products were conducted for TA clone detection (Sangon, Shanghai).

### Immunization

For KLH and NP-KLH immunization, 6- to 8-week-old *STAT1*^mut/WT^ mice and their wild-type controls were immunized with KLH or NP-KLH antigen (1 mg/mL) emulsified in CFA (2 mg/mL, Thermo Scientific) intraperitoneally (200 μL per mouse) or subcutaneously (100 μL per mouse) in the base of the tail. These mice were killed and analyzed individually after 3 days, 7 days or 14 days.

### Cell sorting, culture and stimulation

Naïve CD4^+^T cells were negatively isolated *ex vivo* from spleens and peripheral LNs using Naive CD4^+^T Cell Isolation Kit (Miltenyi Biotec) according to the manufacturer’s protocol, and cultured in flat-bottom 96-well plates previously coated with 1 μg/ml anti-CD3 and 1 μg/ml anti-CD28 in RPMI-1640 medium supplemented with 10% FBS, 2mM L-glutamine, 10mM HEPES buffer solution, 1mM sodium pyruvate, 100 μM nonessential amino acid solution, 50 μM β-mercaptoethanol, 100 U/ml penicillin and 100 μg/ml streptomycin at a density of 1 × 10^5^ cells/well.

For differentiation, naïve CD4^+^T cells were cultured for 3 days with the following cytokines and antibodies for polarization: Th0: anti-IFN-γ (5 µg/mL), anti-IL-4 (5 µg/mL), IL-2 (30 U/mL); Th1: anti-IL-4 (2.5 µg/mL), IL-12 (20 ng/mL), IL-2 (30 U/mL); Th2: IL-2 (30 U/mL), anti-IFN-γ(5 µg/mL), IL-4 (20 ng/mL); Th17: anti-IFN-γ(5 µg/mL), anti-IL-4 (5 µg/mL), IL-6 (30 ng/mL), IL-23 (20 ng/mL), IL-1β(20 ng/mL), TGF-β (2.5 ng/mL); iTreg: anti-IFN-γ(5 µg/mL), anti-IL-4 (5 µg/mL), IL-2 (30 U/mL), TGF-β(5 ng/mL). After differentiation, cells were collected for subsequent experiments. For proliferation, naïve CD4^+^T cells were stained with CellTrace™ CFSE and then stimulated with anti-CD3 and anti-CD28. 72h later, cells were collected for further analysis. For survival, Naïve CD4^+^T cells were isolated and cultured in complete medium with or without IL-7 (10 ng/mL) at 37□ in 5% CO_2_ for indicated time.

CD4^+^T cells were positively isolated from spleens and peripheral lymph nodes using CD4 (L3T4) MicroBeads (Miltenyi Biotec) according to the manufacturer’s protocol. The enriched CD4^+^T cells were stained with CD4, CD44, CXCR5 and PD-1 antibodies for Tfh and non_Tfh sorting and sorted on a BD FACS Aria II cell sorter (BD biosciences). B cells were isolated on a BD FACS Aria II cell sorter by stained with B220 and CD4 antibodies (gated in B220^+^CD4^-^).

### Adoptive transfer and immunization

Naïve CD4^+^T cells from spleens and lymph nodes of CD45.2 STAT1^Mut/WT^ OT-II and CD45.2 OT-II mice were isolated as described in the previous section. 5×10^5^ naïve OT-II cells suspended in PBS were intravenously injected into congenic CD45.1 mice before subcutaneous immunization with 100 μg OVA protein (Sigma-Aldrich) emulsified in CFA (Thermo Scientific). Draining lymph nodes (dLNs) were collected on indicated time post-immunization for subsequent analysis.

For Rag1^-/-^ transfer experiments, B220^+^CD138^-^ plasmacyte depleted B cells, CD4^+^CD25^-^CD44^+^ CXCR5^-^PD1^-^ activated non-TFH cells, and CD4^+^CD25^-^CD44^+^CXCR5^+^PD-1^+^ TFH cells were sorted from WT and *STAT1*^Mut/WT^ mice using a FACSAria III (BD Bioscience). A total of 2×10^6^ B cells and 3×10^5^ non-TFH cells or TFH cells were intravenously transferred into Rag1^-/-^ mice. Experiments were performed 8 weeks after transfer.

### BM chimeras

Bone marrow (BM) chimeric mice were generated by lethally irradiating the indicated recipient mice with 650 Rads, and reconstituted via intravenous (i.v.) injection with1×10^7^ cells of the indicated mix of BM cells. Chimeric mice were maintained on antibiotics for 4 weeks after reconstitution and experiments were performed 10 weeks after reconstitution (2 weeks after KLH and CFA immunization).

### *In vitro* T & B cells co-culture

As previously described (*33*), CD4^+^CD25^-^ CD44^+^CXCR5^-^PD1^-^ activated non-TFH cells, and CD4^+^CD25^-^CD44^+^CXCR5^+^PD-1^+^ Tfh cells were sorted using a FACSAria III (BD Biosciences) from indicated mice. Splenic B220^+^ cells were enriched with B cell isolation kit (Miltenyi Biotec). 3×10^4^ non-Tfh or Tfh cells and 5×10^4^ B220^+^ B cells, or 8 x10^4^ B cells alone were cultured together in 96 U-bottomed plates (Thermo Scientific) pre-coated with 15µg/mL anti-mouse IgM (Jackson Immuno) and 2 µg/mL anti-mouse CD3 (BioXcell) in complete medium at 37□ in 5% CO_2_. After 5 days of co-culture in complete medium, cells and supernatants were harvested for subsequent analysis

### Ruxolitinib treatment

8-week-old *STAT1*^mut/WT^ mice were treated with ruxolitinib (50mg/kg per day, MCE) by oral gavage once daily for 2 weeks. Ruxolitinib was dissolved in 30% polyethylene glycol 300 (PEG300) and diluted with sterile saline to the working concentration. Control mice received vehicle alone.

### IFN-**γ** antibody treatment

6-week-old *STAT1*^mut/WT^ mice were intraperitoneally administrated with 50mg/kg anti-IFN-γ monoclonal antibody (BioXcell, clone: XMG 1.2) or 50mg/kg rat IgG1 isotype control (BioXcell) every 3 days for a total of 7 doses over 3 weeks. Antibodies were diluted in sterile PBS.

### Flow cytometry

For surface staining, *in vitro*-cultured cells or single-cells suspensions isolated from the indicated lymphoid organs were stained for 30min at 4 °C in PBS containing 2% fetal bovine serum (FBS) with fluorochrome conjugated antibodies. Specifically, surface CXCR5 expression was stained using biotinylated anti-CXCR5 (1:100; 2G8; BD Pharmingen) for 1 h at 4□, followed by APC-anti-rat IgG labeled streptavidin (1:200; Jackson Immunoresearch) for 30min at 4□. For intracellular staining, stained cells were fixed and permeabilized with a Cytofix/Cytoperm kit (BD Biosciences), and intranuclear staining was conducted with a Foxp3/Fixation/Permeabilization kit according to the manufacturer’s instructions (eBiosciences). For cytokine staining, cells were incubated in Complete RPMI supplemented with 100 ng/mL Phorbol 12-myristate 13-acetate (PMA) and1 µg/mL ionomycin with 1μg/mL brefeldin A and 2uM monensin to inhibit ER and Golgi transport at 37°C with 5% CO2 for 4 hours. Dead cells were excluded using fixable viability dye staining. For detection of total and phosphorylated STAT1, cells were pre-stimulated by IFN-γ (1 µg/mL) or IFN-α (1 µg/mL) at 37□ for 15 to 30 min, followed by fixation with Phos-flow Fix Buffer I at 37□ for 15 min and permeabilization with BD Phosflow Perm Buffer III at 4□ for 30 min. Cell acquisition was collected on LSR Fortessa or BD Celesta flow cytometer (BD Biosciences) and data was analyzed with FlowJo software (TreeStar, USA). Flow cytometry antibodies are detailed in data file S1.

### ELISA

Serum from immunized mice were collected, and antigen-specific IgM, IgG, IgG1, IgG2a, IgG2b and IgG3 antibodies were measured by using ELISA as previously described(*50, 56*). In brief, Nunc MicroWell 96-well plate was pre-coated with antigens (OVA, NP30-BSA, NP4-BSA, 10 µg/mL) at 4□ overnight. After washing with phosphate buffer saline (PBS) containing 0.05% Tween-20, wells were blocked with PBS containing 1% BSA plus 0.05% Tween-20 at RT for 2 h. The plate was then washed and incubated with 50µl of diluted serum at 4□ overnight. Following incubation, the plate was washed again and incubated with HRP-conjugated secondary antibodies (HRP-IgG, IgG1, IgG2a, IgG2b, IgG2c, IgG3 and IgM, 1:3000) in RT for 1 h. The color reaction was developed using OPD substrate (Sigma-Aldrich) and stopped by adding 1M H2SO4.The plates absorbance was measured at 450nm using a BioTek Synergy H1 plate reader. Alternatively, serum dsDNA autoantibody was measured by ELISA (Alpha Diagnostic) according to the manufacturer’s instructions.

### ANA and dsDNA autoantibody detection via immunofluorescence

The serum ANA and dsDNA autoantibodies were detected using commercial kits (Euroimmun medical diagnostics Co. Ltd) according to the manufacturer’s protocols. Images were captured using an ECLIPSE Ni-E microscope.

### Hematoxylin and Eosin (H&E) staining

The liver, spleen, and lungs from sacrificed mice were fixed in 4% paraformaldehyde and embedded in paraffin. Five-micron sections were stained with hematoxylin & eosin. Images were acquired under a light microscope for histological evaluation.

### Immunofluorescence and immunohistochemistry

For immunofluorescence and confocal imaging, spleen or dLNs were fixed with 1% paraformaldehyde at RT for 30min, followed by incubation in 1% PFA containing 30% sucrose at 4□ overnight, snap-frozen in O.C.T. compound, and then sectioned to 8µm. The tissue sections were stained overnight at 4□ in a humid chamber with the following antibodies: anti-mouse IgD FITC (11-26c.2a, Biolegend), anti-mouse CD4 (17A2, Biolegend), anti-mouse GL7 (GL7, Thermo fisher).

Slides were mounted with permount mounting medium (SP15-100, Fisher scientific). The images were captured using Olympus VS200 and analyzed using Olyvia software.

For immunohistochemistry, paraffin-embedded mouse kidney sections (4 µm) were deparaffinized, rehydrated, and subjected to antigen retrieval in EDTA buffer (pH 8.0) by microwave heating. Endogenous peroxidase was quenched with 3% H□O□, followed by blocking with 3% BSA. Sections were incubated with primary antibodies, then with HRP-conjugated secondary antibodies (1:10,000). Signal development was achieved using DAB, followed by hematoxylin counterstaining, dehydration, and mounting with neutral resin.

### Western blot

Detection of pSTAT1 and total STAT1 was performed as described previously(*44, 88*). Briefly, splenic single-cell suspension, PBMCs or U3C cell line (transfected with WT and mutant plasmids, gifted by Satoshi Okada) were obtained. Cells were stimulated with IFN-γ (1 µg/mL) at 37□ for 30 min and subsequently lysed in RIPA buffer (150mM NaCl, 1% NP-40, 0.5% sodium deoxycholate, 0.1% sodium dodecyl sulfate, 50mM Tris, 1mM EDTA and 1mM EGTA) supplemented with 2 mM phenylmethylsulfonylfluoride (PMSF) and protease inhibitor cocktails (Roche) on ice for 40 min. Proteins were separated and transferred to PVDF membranes (Millipore). Membranes were incubated with HRP-conjugated β-actin monoclonal antibody (1:5000), phospho-STAT1 (Tyr701) (1:1000) or STAT1(1:1000) monoclonal antibody followed by HRP-conjugated goat anti-rabbit IgG secondary antibody (1:5000) at 4 °C overnight. Following three consecutive 10-minute washes with TBST, the blots were incubated with the appropriate horseradish peroxidase-conjugated secondary antibody (1:5000, Cell Signaling Technology, USA) for 1 h at room temperature. The blots were scanned using the Odyssey Infrared Imaging System (LI-COR Biosciences, USA). Images were analyzed using ImageJ software.

### Quantitative PCR

Total RNA was extracted with Trizol reagent, and first strand cDNA was synthesized using PrimeScript RT Master Mix. Quantitative real-time PCR was conducted with a TB Green® Premix Ex Taq™ kit (Tli RNaseH Plus) (Takara Bio, China) and a CFX96 real-time system PCR system (Bio-Rad, USA) according to the manufacturers’ instructions. Gene expression levels were analyzed with the comparative Ct method and normalized to β-actin housekeeping gene levels. Primers were listed in data file S2.

### RNA library preparation, sequencing and analysis

Total RNA was extracted with the TRIzol reagent according to the manufacturer’s instructions. For library preparation, 500ng of total RNA was processed using the NEBNext® Ultra™ RNA Library Prep Kit for Illumina® (NEB, USA), following the manufacturer’s protocol. Briefly, mRNA was enriched using poly-T oligo-attached magnetic beads, then fragmented with divalent cations at elevated temperatures. First-strand cDNA synthesis was performed using random hexamer primers and M-MuLV Reverse Transcriptase (RNase H), followed by second-strand synthesis with DNA Polymerase I and RNase H. After end repair and 3’ adenylation, NEBNext adaptors with a hairpin loop structure were ligated. Size selection (250–300 bp) was performed using the AMPure XP system (Beckman Coulter). The adaptor-ligated cDNA was treated with USER Enzyme (NEB, USA), followed by PCR amplification using Phusion High-Fidelity DNA polymerase and indexed primers. Final libraries were purified and quality-checked with the Agilent Bioanalyzer 2100. Libraries were clustered using the cBot system and sequenced on an Illumina NovaSeq platform with 150 bp paired-end reads.

Raw sequencing data were subjected to quality control using FastQC (v0.12.1). FASTP (v0.23.4) was employed to remove sequencing adapters and low-quality bases, retaining bases with a Phred quality score ≥ 20 and using the Phred33 quality encoding system. Filtered reads were aligned to the mm10 mouse reference genome using HISAT2(v2.2.1) with default parameters. Gene expression quantification was performed using the feature Counts tool from the Subread package, which was then normalized using Transcripts Per Kilobase of exon model per Million mapped reads (TPM). Finally, TPM values were used to evaluate gene expression levels. Differential gene expression analysis was conducted using DESeq2. Differentially expressed genes (DEGs) were identified with a fold change (FC) ≥ 1.5 and a false discovery rate (FDR)-adjusted p-value < 0.05. Additional downstream analyses and visualizations—including principal component analysis (PCA) plots, clustered heatmaps, and volcano plots—were performed using R. Comparison of our data with published datasets or our in-stock data was accomplished using Gene Set Enrichment Analysis (GSEA). Genes for GSEA were ranked by -log10(p-value) times the sign of the fold change for the indicated comparisons.

### CUT & Tag library preparation, sequencing and analysis

Mice CD4^+^CD25^-^CD44^+^CXCR5^-^PD1^-^ non-TFH cells and CD4^+^CD25^-^CD44^+^CXCR5^+^PD-1^+^ TFH cells were sorted using a FACSAria III (BD Biosciences), and then stimulated with IFN-γ for 15min. Following stimulation, Library from indicated group was prepared using Hyperactive Universal CUT&Tag Assay Kit for Illumina according to protocol. Subsequently, pair-end sequencing of sample was performed on Illumina platform (Illumina, CA, USA). Library quality was assessed on the Agilent Bioanalyzer 2100 system by Knorigene Technologies (Chongqing, China).

Raw CUT & Tag FASTQ data were processed with FASTP (v0.23.4) to remove adapter sequences, poly-N reads, and low-quality reads. Q20, Q30 and GC content were calculated, and all downstream analyses used high-quality clean reads. Reference genome (with annotation files from ENSEMBL) was indexed via BWA (v0.7.17); clean reads were aligned to the genome using BWA mem. Aligned SAM files were converted to BAM format, sorted by genomic coordinates, and further processed with alignmentSieve (v3.5.5) for filtering: PCR duplicates and reads mapping to blacklist regions were removed, with additional parameters-minMappingQuality 25 and -samFlagExclude 260. Peak calling was performed using MACS3 (v3.0.0) with a q-value threshold of 0.05. Peak distribution, width, enrichment, significance and summit counts were analyzed. Homer was used for de novo motif discovery and known motif identification. Peaks were annotated by ChIPseeker R package (v1.44.0), followed downstream analysis using the R package clusterProfiler (v4.16.0). Differential peak analysis was performed using the DiffBind R package (v3.18.0). Differential peaks were identified based on fold change in peak enrichment with an absolute value ≥ 2 and adjusted *p*-value < 0.05, data were viewed and presented with the Integrative Genomics Viewer (IGV).

### Statistical Analysis

Unless indicated otherwise, statistical analyses were performed using the two tailed unpaired or paired Student’s *t* test with Graphpad Prism 10.0. All data are shown as means ± SEM. Differences were considered statistically significant at *P* < 0.05.

## RESULTS

### STAT1-GOF Patients Presented with Autoimmune Manifestations and Displayed an Increased Abundance of Circulating Tfh Cells

We studied 20 patients with germline *STAT1*-GOF mutations by whole-exome sequencing (Fig. 1A). Of these, seventeen distinct heterozygous *STAT1* mutations were identified. Six patients (P6, P7 and P13 to P16) have not been previously described (Fig. S1A). Two patients, P6 and P7, harbored novel deletion mutations: p.L166_D168del and p.H386_T387del, respectively (Fig. S1A-B), which showed enhanced expression of total STAT1 and phosphorylated STAT1 (Y701, pSTAT1) upon IFN-γ or IFN-α stimulation (Fig. S1C-D). Transient expression experiments further confirmed that p.L166_D168del, p.H386_T387del, as well as the known GOF mutant p.R274Q, resulted in increased levels of total STAT1 and pSTAT1 compared to control (Fig. S1E).

**Fig. 1.**
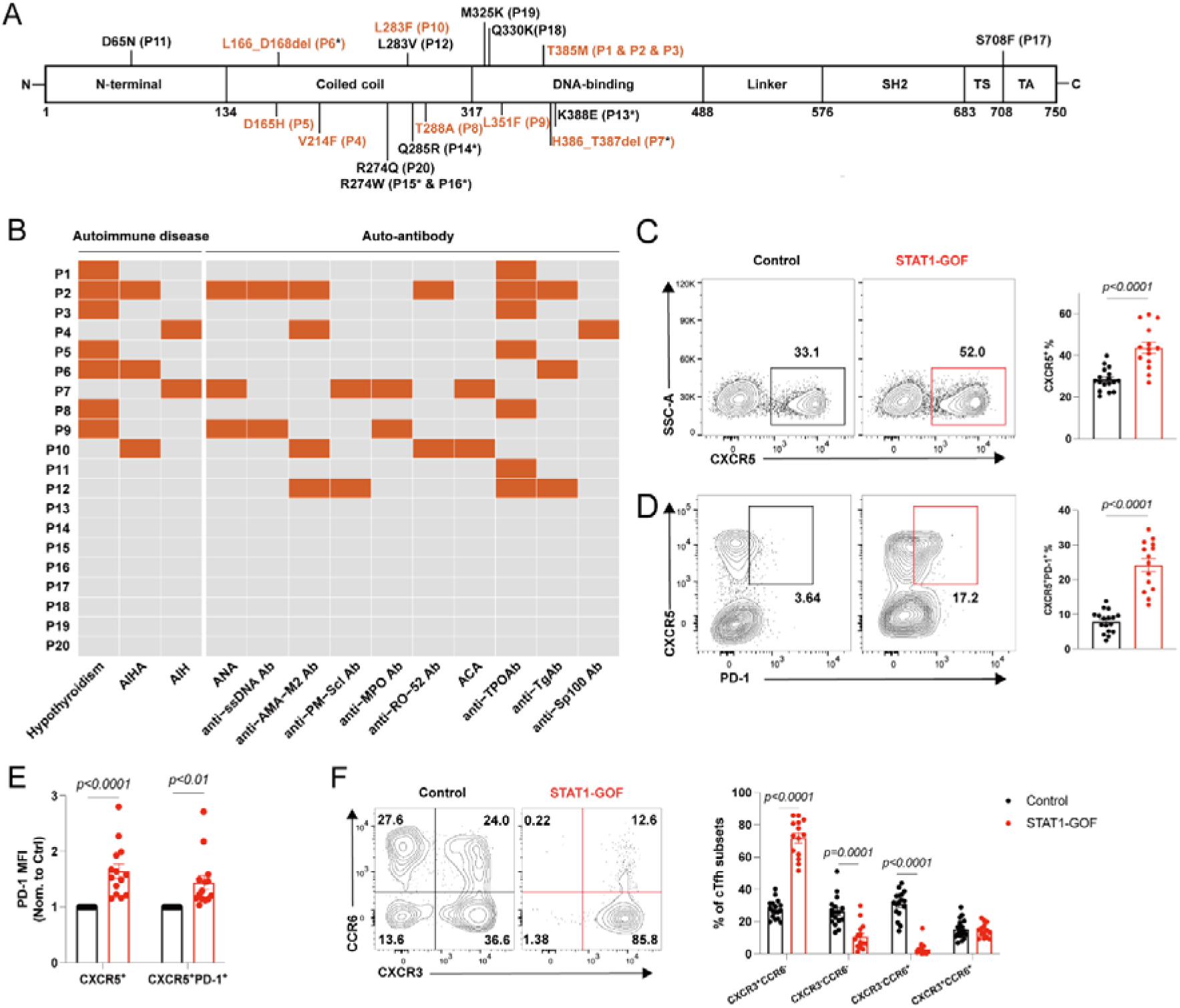
*STAT1*-GOF patients exhibit autoimmune manifestations and display a skewed cTfh compartment. (**A**) Sixteen pathogenic mutations in STAT1 coding region carried by 20 patients are indicated. Red text denotes patients with autoimmune phenotypes. The asterisks (*) indicate newly identified patients. (**B**) Heatmap illustrating autoimmune phenotypes in patients. Red and grey shading represents positive and negative status, respectively. (**C**) Representative flow cytometry plots and pooled data of the percentage of circulating CXCR5^+^ T cells in CD3^+^CD4^+^CD45RO^+^ T cells. (**D**) Representative flow cytometry plots and pooled data of the percentage of circulating CXCR5^+^PD-1^+^ in CD3^+^CD4^+^CD45RO^+^ T cells. (**E**) Bar plots showing PD-1 MFI (normalized to aged-matched controls). (**F**) Representative flow cytometry plots and pooled data of the percentage of cTfh subsets in circulating CXCR5^+^CD45RA^-^CD4^+^T cells. MFI: Mean fluorescence intensity. cTfh: circulating Tfh. Numbers adjacent to outlined areas or in the indicated quadrants represent the percentage of cells in the area. Each symbol represents an individual throughout. Data were represented as means ± SE and analyzed with two-tailed unpaired *t* test.

In our cohort, apart from developed CMC caused by fungal pathogens (n=18), patients with *STAT1-*GOF mutations showed susceptibility various bacterial and viral infections (table S1). Notably, 50% of patients exhibited clinical autoimmune manifestations, such as hypothyroidism (n=7), autoimmune hemolytic anemia (n=3), autoimmune hepatitis (n=2), and two additional patients (P11 and P12) tested positive for autoantibodies despite having no clinical signs of autoimmunity at the time of assessment (Fig. 1B, table S1).

Given these frequent autoimmune manifestations, we evaluated the cTfh cells in the peripheral blood of patients with *STAT1-*GOF mutations. Despite the similar proportion of activated CD45RA^-^ CD4^+^T cells between the *STAT1-*GOF patients and age-matched healthy control subjects (Fig. S1F), the patients exhibited an obviously increased in the abundance of both CD4^+^CD45RA^-^CXCR5^+^ cTfh cells and CD4^+^CD45RA^-^CXCR5^+^ PD-1^+^cTfh cells (Fig.1C-D). The cTfh cells from the patients also displayed elevated expression of PD-1(Fig. 1E), which is correlated with active Tfh cell differentiation and the amounts of autoantibodies (*21*). Moreover, consistent with others reports(*37, 47, 48*), cTfh in STAT1-GOF patients were in favor of CXCR3^+^ Tfh1-like differentiation and deviated from CCR6^+^Tfh17-like phenotype differentiation (Fig.1F). Thus, the biasedly increased cTfh in *STAT1*-GOF patients may be closely associated with the pathophysiological development of autoimmunity.

### Spontaneous Development of Autoimmunity in *Stat1*-GOF Mice

To explore the effect of STAT1 GOF on immune responses, we generated a mouse model that the murine *Stat1* exon 14 was substituted to introduce the T385M mutation *in situ*, which corresponds to the common gain-of-function mutation site in *STAT1*-GOF patients (T385M) (Fig. S2A-B).

Heterozygous and homozygous mutant mice (hereafter referred to as *Stat1*^Mut/WT^, *Stat1*^Mut/Mut^ mice) were fertile and born at normal Mendelian ratios and did not show obvious behavioral defects.

Splenocytes from mutant mice show increased levels of total STAT1 and IFN-γ-induced pSTAT1 compared to wide-type (WT) control (Fig. S2C-D). Dephosphorylation kinetics of pSTAT1 was delayed in mutant mice following IFN-γ treatment (Fig. S2C-D). Also, the expression of total STAT1 and IFN-γ-stimulated pSTAT1 were elevated in genetic dose-dependent manner in CD4^+^ T and B220^+^ cells (Fig. 2A-B). Thus, the *Stat1* T385M mutant can serve as a *Stat1* GOF mice model.

**Fig. 2.**
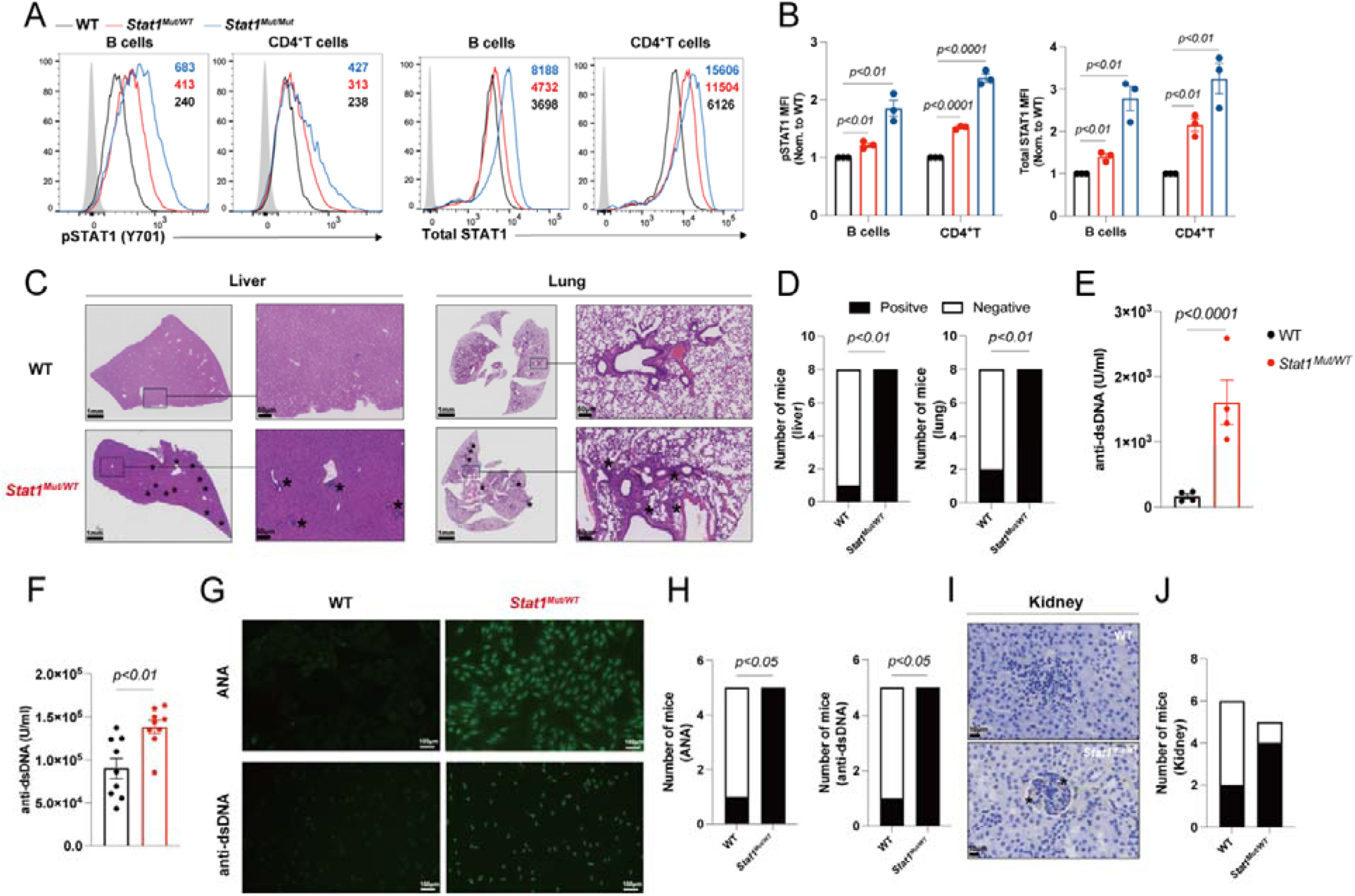
*Stat1*-GOF mice spontaneously develop autoimmune phenotype. (**A**) Histogram of pSTAT1 (Y701) and total STAT1 in B cells and CD4^+^T cells. (**B**) pSTAT1 and total STAT1 MFI in indicated subsets. (**C**) H&E staining of liver sections (left) and lung sections (right). (**D**) Bar graph showing numbers of mice with positive or negative inflammatory cell infiltration in indicated tissue. (**E-F**) The levels of anti-dsDNA antibody detected in serum in mice at 6-8 weeks (E) and 20-22 (F) weeks. (**G**) Representative Hep-2 ANA (up) and *Crithidia luciliae* anti-dsDNA antibody (down) staining. (**H**) Bar graph showing numbers of mice with positive or negative results in indicated autoantibody. (**I**) Representative staining of IgG in kidney. (**J**) Bar graph showing numbers of mice with positive or negative IgG deposition in kidney. Each symbol represents an individual mouse throughout. Data are representative of three independent experiments. Data were represented as means ± SEM and analyzed with two-tailed unpaired *t* test.

*Stat1* GOF mice displayed splenomegaly with increased splenic cellularity, lymphadenopathy (Fig. S2E-F), and massive lymphocyte infiltration of the bronchioles and blood vessels of the lung, and scattered lymphocyte hyperplasia in the liver compared to control mice (Fig. 2C-D). At 6 weeks of age, *Stat1*^Mut/WT^ mice already had more anti-dsDNA IgG than littermate controls (Fig. 2E). With age, by 20 weeks, higher levels of anti-dsDNA IgG and anti-ANA IgG autoantibodies were positively detected in the serum of all *Stat1*^Mut/WT^ mice (Fig. 2F-H), and IgG immune complexes causing tissue damage were deposit at glomerulus of mutant mice (Fig. 2I-J). Overall, these findings suggest that *Stat1* ^Mut/WT^ mice exhibited multiorgan inflammation and spontaneous autoimmune phenotype, and that the mutant mouse model can at least partly recapitulate the pathogenesis of *STAT1*-GOF patients.

### Increased Numbers of Tfh Cells and Spontaneous GC Formation in *Stat1*-GOF Mice

To evaluate how STAT1 GOF leads to autoimmunity, we analyzed T cell and B cell populations. WT and mutant mice had comparable numbers of total thymocytes, except for slight increase in the proportions of single-positive CD4^+^ and CD8^+^ T cells in the *Stat1* ^Mut/WT^ mice (Fig. S3A-B). In the spleen, *Stat1* ^Mut/WT^ mice had fewer percentages of T cells and an altered ratio of CD4^+^ cells to CD8^+^ cells relative to that of their wild-type littermates (Fig. S4A-D). Among the conventional T cell compartments, *Stat1* ^Mut/WT^ mice had a significantly lower percentage of CD62L^hi^CD44^lo^ naïve T cells and a correspondingly expanded proportion of CD44^hi^ cells with an activated or memory phenotype in the spleen and peripheral lymph nodes (LNs) than in that of WT control (Fig. S4E-H). This alteration was further exacerbated over time (Fig. S5A-E). These results suggested that STAT1 GOF disrupted the homeostasis of peripheral T cells in mice and promoted the accumulation of activated T cells.

Similar to results obtained for recent reports(*47–49*), the proportion and numbers of CD4^+^ CD44^+^CXCR5^+^ PD-1^+^ FoxP3^-^Tfh cells were significantly higher in the spleen and peripheral LNs of *Stat1* ^Mut/WT^ mice compared to their littermate controls(Fig. 3A, Fig. S6A). Bcl-6^hi^ GC Tfh cell populations were also expanded in *Stat1* ^Mut/WT^ mice (Fig. 3B, Fig. S6B). In addition, Tfh cells from mutant mice expressed high levels of key Tfh cell markers, such as CXCR5, PD-1 and Bcl-6 (Fig. 3C). In contrast, the ratio of Tfh cells to Foxp3^+^ follicular helper regulatory T cells (Tfr cells) in the *Stat1* ^Mut/WT^ mice remained similar to that of controls (Fig. 3D).

**Fig. 3.**
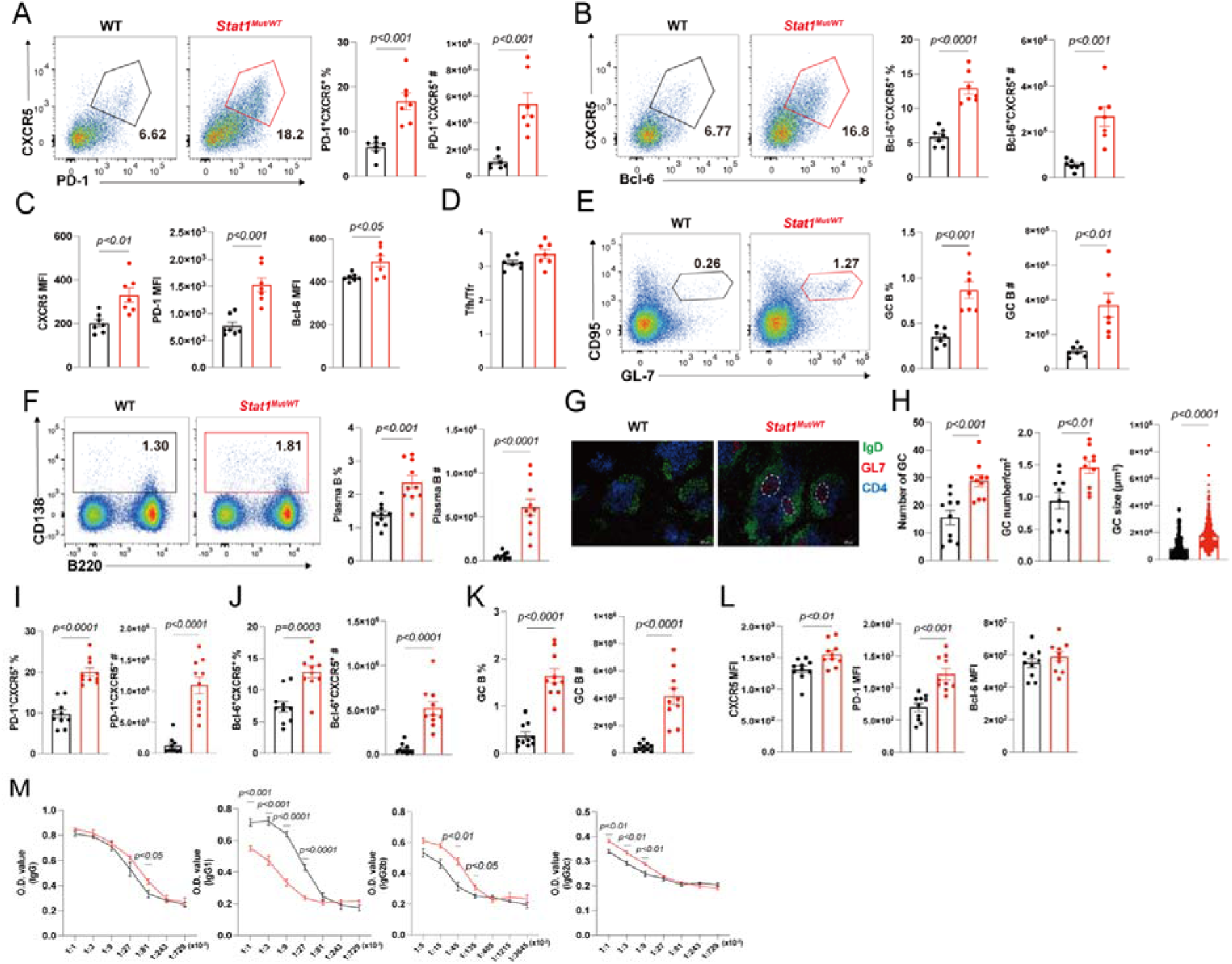
STAT1 GOF mice exhibit spontaneous expansion of Tfh and GC B cells. (**A-B**) Representative flow cytometry plots and pooled data of the percentage and numbers of splenic CXCR5^+^ PD-1^+^Tfh cells (A) and CXCR5^+^Bcl-6^+^ Tfh cells(B) in CD4^+^CD44^+^Foxp3^-^T cells. (**C**) MFI of CXCR5, PD-1 and Bcl-6 in CD4^+^CD44^+^Foxp3^-^T cells. (**D**) The ratio of Tfh (CD4^+^CD44^+^Foxp3^-^Bcl-6^+^CXCR5^+^) to Tfr (CD4^+^CD44^+^Foxp3^+^Bcl-6^+^CXCR5^+^) percentages. (**E-F**) Representative flow cytometry plots and pooled data of the percentage and numbers of splenic GC B cells (E) and plasma B cells (F). (**G**) Representative immunofluorescence images of spleen sections show GC (outlined by the white dashed line), featured by IgD (green), GL7 (red) and CD4 (blue). (**H**) Bar graphs showing GC number, GC number per cm^2^ and GC size. (**I-K**) Percentage and number of splenic CXCR5^+^PD-1^+^ Tfh (I), CXCR5^+^Bcl-6^+^ Tfh (J) and GC B cells (K) in mice at 20-22 weeks. (**L**) MFI of CXCR5, PD-1 and Bcl-6 in CD4^+^CD44^+^Foxp3^-^T cells from mice at 20-22 weeks. (**M**) Serum total IgG and IgG subtype levels in mice. Numbers adjacent to outlined areas represent the percentage of cells in the area. Each symbol represents an individual mouse throughout. Data are representative of three independent experiments. Data were represented as means ± SEM and analyzed with two-tailed unpaired *t* test.

In parallel, the population of B220^+^GL7^+^CD95^+^ GC B cells in both the LNs and the spleen, as well as more CD138^+^ plasma cells, were significantly increased in *Stat1* ^Mut/WT^ mice than in wild-type mice (Fig. 3E-F, Fig. S6C). In addition, substantially more GCs were spontaneously formed in the spleens of mutant mice (Fig. 3G-H). In consistent with time-dependent lymphocyte activation, the elevated Tfh and GC B cells more visible in mutant mice with age (Fig. 3I-L). To dissect contribution of Tfh to the exaggerated GC response, B cells were co-transferred with sorted Tfh from either WT or *Stat1* ^Mut/WT^ mice into Rag1^-/-^ recipients, respectively. 8 weeks after reconstitution, mice receiving *Stat1* ^Mut/WT^ Tfh together with WT B cells exhibited highly elevated serum anti-dsDNA antibody titers and a markedly increased frequency of GC B cells, compared with recipients reconstituted with WT Tfh and WT B cells (Fig. S7A-B), indicating that *Stat1* ^Mut/WT^ Tfh cells are sufficient to exacerbate GC response and promote autoantibody production.

Furthermore, we detected the levels of serum immunoglobulin, and observed that the concentrations of total IgG, IgG2b and IgG2c were higher in *Stat1* ^Mut/WT^ mice than controls, whereas the concentrations of IgG1 were much lower than controls (Fig. 3M), indicating that circulating immunoglobulin in mutant mice was skewed primarily toward IgG2 subclass, typically associated with type 1 immune response. These data suggest that enhanced Tfh cell and spontaneous GC responses likely drive the development of autoimmunity in STAT1-GOF patients.

### Hyperactivated STAT1 intrinsically promotes the Tfh differentiation

To determine the effects of STAT1 GOF on development of Tfh cells, we firstly immunized mice intraperitoneally with T cell-dependent antigen keyhole limpet hemocyanin (KLH) emulsified in Complete Freund’s Adjuvant (CFA). As expected, on day 14 post-immunization, frequencies and numbers of both CD4^+^CXCR5^+^PD-1^+^ and CD4^+^CXCR5^+^Bcl-6^+^ Tfh cells were significantly increased in *Stat1* ^Mut/WT^ mice compared with WT mice (Fig. 4A-C). The elevation in the Tfh cell population in *Stat1* ^Mut/WT^ mice was probably not due to altered proliferation or apoptosis (Fig. S8A-B). Similarly, following immunized with NP-KLH emulsified in CFA, there was also a greater frequency of Tfh cells than that of controls (Fig. 4D-E). Next, we infected *Stat1* ^Mut/WT^ mice and WT mice with LCMV Armstrong. At day 8 after infection, the frequency of CD44^+^CXCR5^+^SLAM^lo^ Tfh and CD4^+^CD44^+^CXCR5^+^Bcl-6^+^ Tfh cells were also increased in mutant mice (Fig. S8C-F). These data indicated that STAT1 GOF promoted Tfh differentiation either in type 1 or type 2 immunization.

**Fig. 4.**
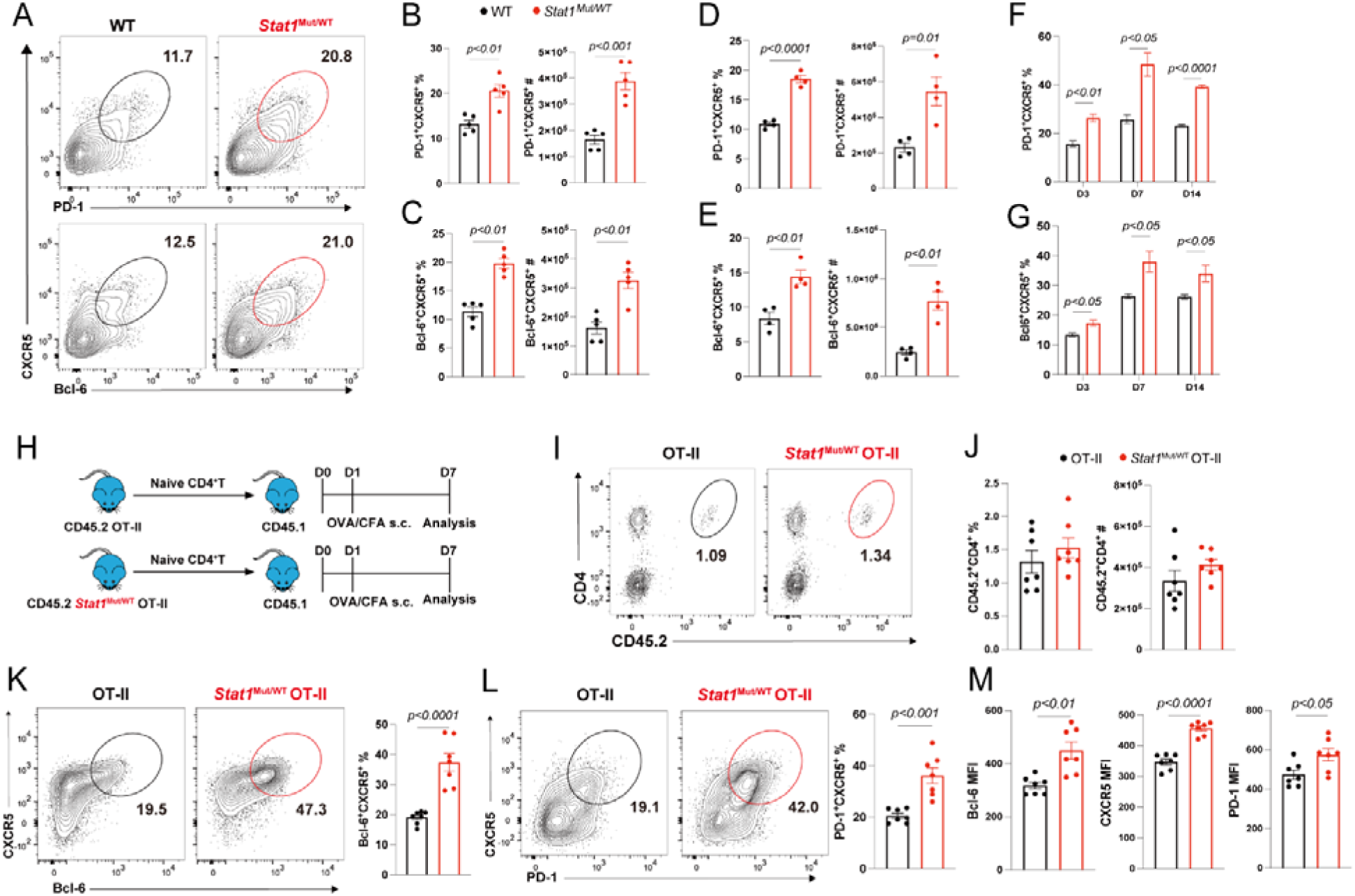
Hyperactivated STAT1 intrinsically promotes the Tfh differentiation. **(A-C)** The indicated mice were intraperitoneally immunized with KLH/CFA. 14 days after the immunization, splenic Tfh cells were determined. Representative flow cytometry plots and pooled data of the percentage and numbers of splenic CXCR5^+^PD-1^+^ and CXCR5^+^Bcl-6^+^ Tfh cells in CD4^+^CD44^+^Foxp3^-^T cells. **(D-E)** The indicated mice were intraperitoneally immunized with NP-KLH/CFA. 14 days after the immunization, the percentages and numbers of splenic CXCR5^+^PD-1^+^ and CXCR5^+^Bcl-6^+^ Tfh cells were determined. **(F-G)** The indicated mice were immunized subcutaneously with KLH/CFA. The percentages of Tfh at indicated time points were determined. **(H-M)** Naïve OT-II cells from indicated mice were isolated and transferred into congenic mice. The recipient mice were subcutaneously immunized with OVA/CFA. Seven days after the immunization, Tfh cells in dLNs were determined. **(H)** Experimental design for evaluating the role of mutant *Stat1* in Tfh differentiation. **(I-J)** Representative flow cytometry plots and pooled data of the percentage and number of donor cells. **(K-L)** Representative flow cytometry plots and pooled data of the percentage of CXCR5^+^PD-1^+^ and CXCR5^+^Bcl-6^+^in donor cells. **(M)** MFI of CXCR5, PD-1 and Bcl-6 in donor cells. Numbers adjacent to outlined areas represent the percentage of cells in the area. Each symbol represents an individual throughout. Data are representative of three independent experiments. Data were represented as means ± SEM and analyzed with two-tailed unpaired *t* test.

To test whether STAT1 GOF is important for the early fate commitment of Tfh cells, we dynamically monitored Tfh cells at different time points after immunization, and found that at day 3 after subcutaneously immunization, the abundance of Tfh was higher in draining lymph nodes (dLNs) in *Stat1* ^Mut/WT^ mice compared with WT mice (Fig. 4F-G). This suggested that overactivated STAT1 promoted early Tfh differentiation *in vivo*.

To further address whether STAT1 GOF intrinsically results in enhanced capacity of naïve T cell differentiation into Tfh cell, we transferred naïve OT-II cells into congenic mice and immunized with OVA plus CFA and analyzed at later time point (Fig. 4H). At day 7 after immunization, we observed significantly more (2-fold) CXCR5^+^PD-1^+^ and CXCR5^+^Bcl-6^+^ Tfh cells in transferred *Stat1* ^Mut/WT^ cells than that in OT-II WT cells (Fig. 4K-L), despite the comparable transferred OT-II cell numbers (Fig. 4I-J). The expressions of CXCR5, PD-1 and Bcl-6 were also increased in transferred mutant OT-II cells compare with control (Fig. 4M). Furthermore, we reconstituted the irradiated Rag1 KO mice with equally mixed bone marrow from *Stat1* ^Mut/WT^ (CD45.2^+^) and WT cells (CD45.1^+^) (Fig. S9A). Following immunization, we observed that CD4^+^ T cell populations derived from *Stat1* ^Mut/WT^ origin showed greater frequency of CXCR5^+^PD-1^+^ Tfh cells than those from WT donor (Fig. S9B). Collectively, these data indicated that hyperactive STAT1 intrinsically promoted Tfh differentiation.

### Defective T Cell-Dependent Humoral Responses in *Stat1*-GOF Mice

To investigate the downstream effect of increased Tfh cells in *Stat1* GOF mice, we analyzed GC responses. We firstly inoculate mice with various antigen and analyzed the antigen-specific antibody production. After KLH immunization, the frequency and numbers of GC B cells were significantly increased in the mutant mice compared to WT control (Fig. 5A). To assess the antigen-specific humoral response, we further treated mice with NP-KLH precipitated in CFA. Although the total GC B cells in mutant mice were significantly expanded (Fig. 5B), the proportion of the antigen-binding (NP^+^) GC B were lower in *Stat1* ^Mut/WT^ mice than that in WT mice (Fig. 5C). This was coupled with a serum antibody isotype shift towards reduced NP-specific IgG1 and elevated NP-specific IgG2a/2c in mutant mice (Fig. 5D-E). Consistently, systemic LCMV *Armstrong* infection led to decreased IgG1^+^ but increased IgG2a^+^ GC B in *Stat1* ^Mut/WT^ mice (Fig. S10A-B). Nonetheless, following NP-KLH immunization, the ratio of high-affinity antibodies to NP4 to antibodies to NP30, which is often used to evaluate antibody affinity maturation, in serum of *Stat1* ^Mut/WT^ mice was similar to WT controls (Fig. S10C). These fundings indicated that STAT1 GOF results in a dysregulated GC response, biasedly inducing GC B toward to IgG2a/c isotype switching.

**Fig. 5.**
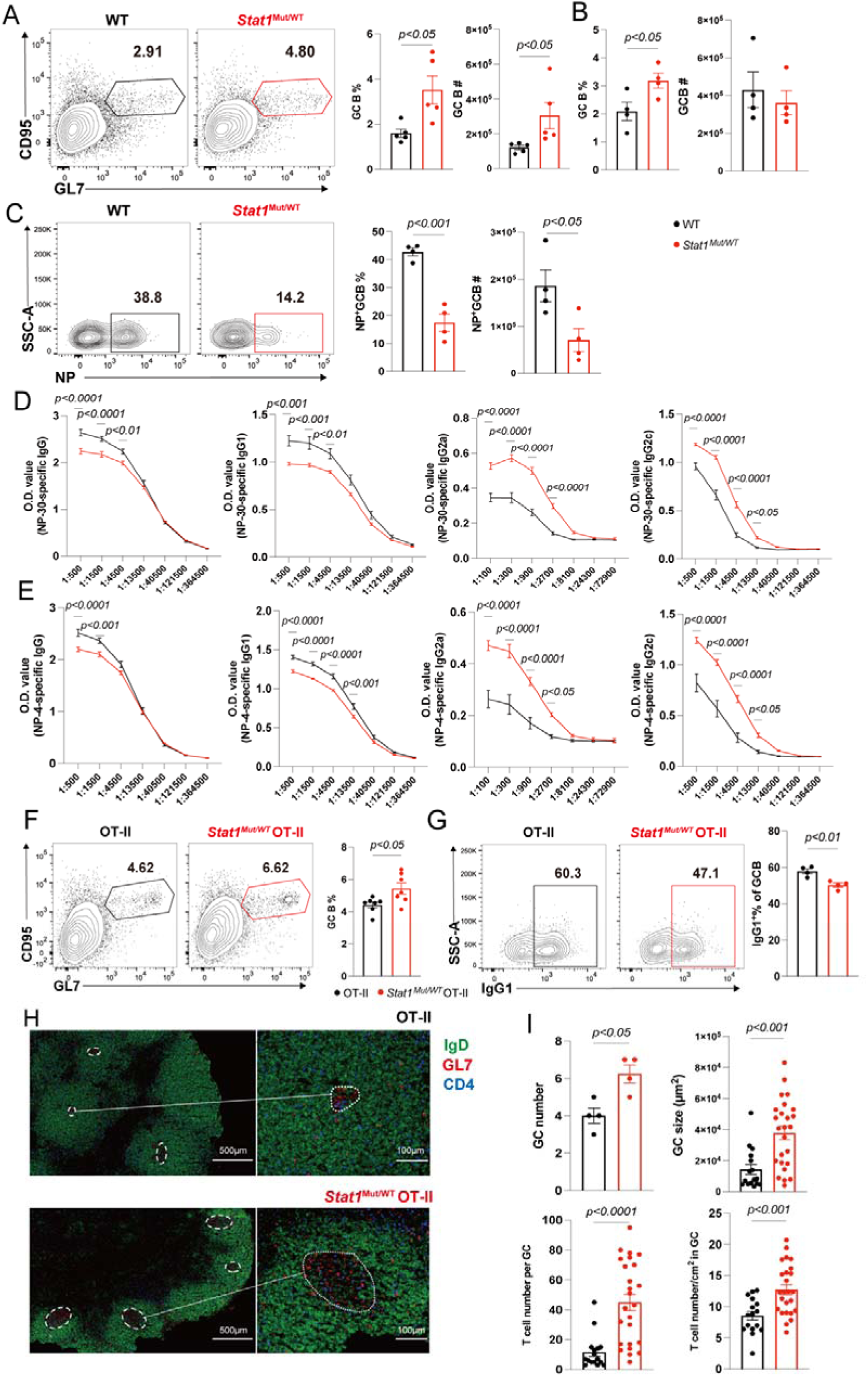
Defective T cell-dependent humoral responses in *Stat1*^Mut/WT^ mice. **(A)** The indicated mice were intraperitoneally immunized with KLH/CFA. 14 days after the immunization, splenic GC B cells were determined. Representative flow cytometry plots and pooled data of the percentage and numbers of splenic GC B cells in B220^+^ B cells. **(B)** Percentage and number of GC B cells after immunized with NP-KLH/CFA. **(C)** Representative flow cytometry plots and pooled data of the percentage and numbers of splenic NP-specific GC B cells. **(D-E)** NP-30-specific and NP-4-specific IgG subtypes in serum. **(F-I)** Naïve OT-II cells from indicated mice were isolated and transferred into congenic mice. The recipient mice were subcutaneously immunized with OVA/CFA. Seven days after the immunization, GC B cells in dLNs were determined. **(F-G)** Representative flow cytometry plots and pooled data of the percentage of GC B cells (F) and IgG1^+^GC B cells (G)in B220^+^ B cells from recipient mice. **(H)** Distribution of donor cells in B-cell follicles from dLNs in congenic mice immunized with OVA/CFA for 7 days. IgD (green), GL7 (red), CD45.2 (blue). **(I)** GC numbers, GC size, donor OT-II cells number in each GC were shown. Numbers adjacent to outlined areas represent the percentage of cells in the area. Each symbol represents an individual throughout. Data are representative of three independent experiments. Data were represented as means ± SEM and analyzed with two-tailed unpaired *t* test.

To examine the function roles of Tfh cells *in vivo*, we then transferred the mutant or WT naïve OT-II cells into congenic mice and immunized with OVA/CFA, which primarily elicits a type 2 response and promotes IgG1 production. At day 7 post immunization, GC B cells, GC B numbers and the size of GC in mice receiving mutant OT-II cells were obviously increased (Fig. 5F and 5H-I). In addition, we found that more mutant OT-II cells accumulated in the GC area in dLNs. However, consistent with impaired class switching, IgG1^+^ GC B was also decreased in mutant transferred mice accompanied by decreased serum anti-OVA IgG1 while elevated anti-OVA IgG2c level (Fig. 5G, Fig. S10D). Thus, echoing previous studies for *STAT1*-GOF patients, *Stat1* ^Mut/WT^ mice exhibited striking alterations in humoral immunity, characterized by increased formation of GCs largely owing to uncontrolled Tfh differentiation, but defects in class-switched antigen-specific antibody responses.

### Acquired Th1 Transcriptomic Features in Tfh cells of *Stat1*-GOF mice

To gain insight into the potential mechanisms in which hyperactive STAT1 promoted Tfh differentiation, we performed RNA-seq analysis on non-Tfh cells and Tfh cells isolated from WT mice and *Stat1* ^Mut/WT^ mice following immunization, respectively (Fig. 6A-B, Fig. S11A). STAT1 is an evolutionally cytoplasmic transcription factor that mediates the signaling of IFN-α/β and IFN-γ, thereby playing an indispensable role in the T cells immune defense against various pathogens.

**Fig. 6.**
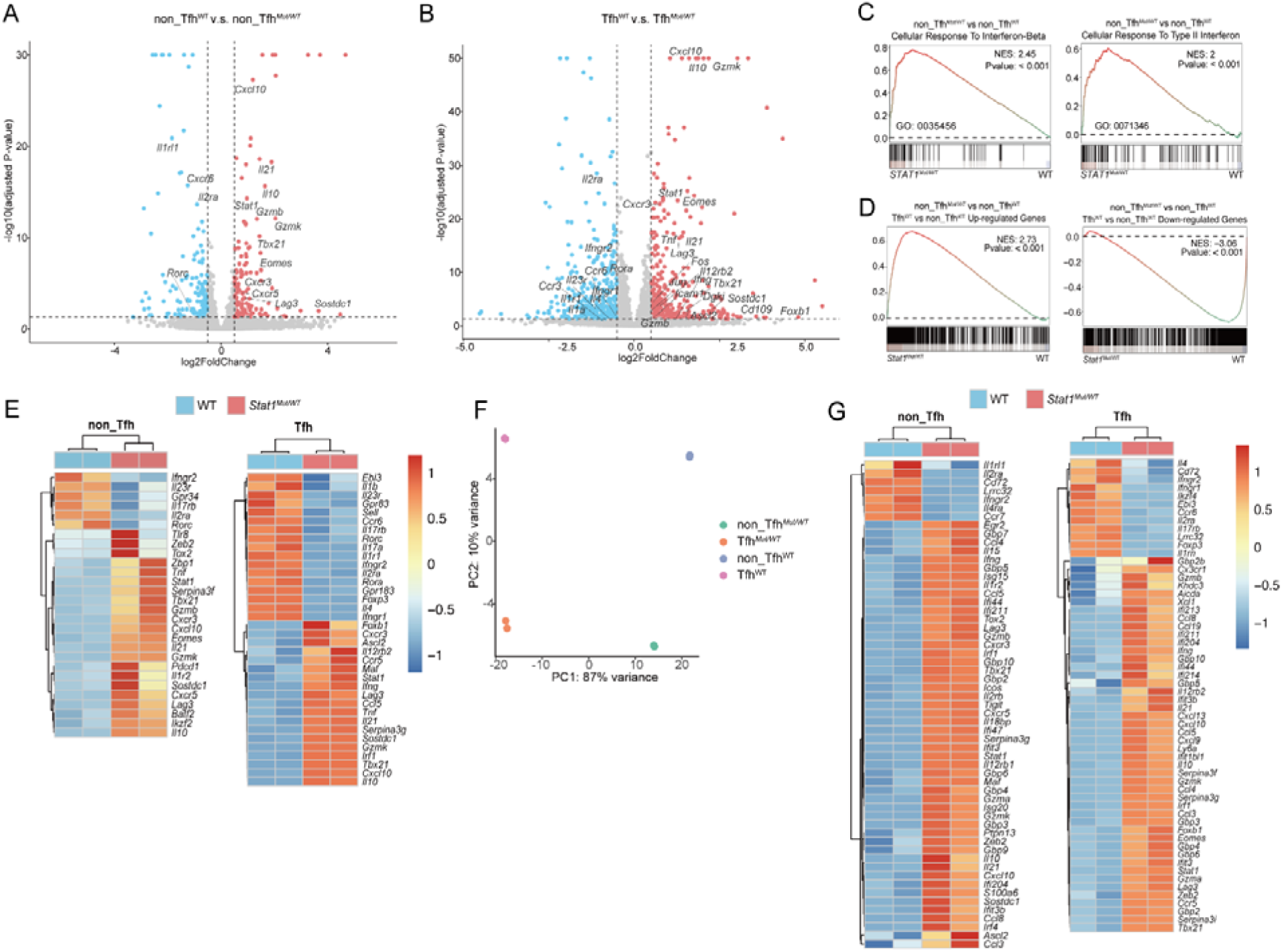
Acquired Th1 transcriptomic features in Tfh cells of *Stat1* ^Mut/WT^ mice. (**A-B)** Volcano plots showing DEGs of non-Tfh (A) and Tfh (B) between WT and *Stat1^Mut/WT^*. Cells were sorted from mice immunized with KLH/CFA for 14 days. **(C-D)** GSEA plot for the indicated gene set. **(E)** Heatmap visualization of DEGs in non-Tfh and Tfh between WT and *Stat1^Mut/WT^* mice immunized with KLH/CFA for 14 days. **(F)** PCA plot of the transcriptomic data of indicated cell groups from mice at age of 20-22 weeks. Each dot represents an individual sample. **(G)** Heatmap visualization of DEGs in non-Tfh and Tfh between WT and *Stat1^Mut/WT^* at age of 20-22 weeks. DEGs: differentially expressed genes. GSEA: gene set enrichment analysis. PCA: principal component analysis.

Consistent with this notion, the differentially expressed genes in non-Tfh cells from mutant mice were positively enriched in signaling pathway for cellular response to interferon-β and cellular response to type II IFN (Fig. 6C). Th1 lineage related genes, such as *Tbx21*, *Eomes*, *Gzmb* and *Tnf*, were increased whereas Th17 related genes (*Rorc, Il23r* and *Il17rb*) were suppressed in mutant non-Tfh cells. Of note, the expression of key regulators of Tfh differentiation was increased in non-Tfh cells from *Stat1* ^Mut/WT^ mice, including *Cxcr5*, *Sostdc1*(*50*), *Pdcd1*, *Tox2*(*51, 52*), *Zeb2* (*53, 54*)and effector cytokines *Il21* (Fig. 6E). Thereby, the differentially expressed 1,441 genes in Tfh cells compared with non-Tfh cells from WT (changed over 2 folds, data file S3) were identified and choose for gene set enrichment analysis (GSEA). This analysis revealed that *Stat1* ^Mut/WT^ non-Tfh cells showed enrichment for the gene set upregulated in Tfh cells, while the gene set restrained in Tfh cells was more enriched in WT non-Tfh cells relative to mutant cells (Fig. 6D). With published data sets (GSE24574)(*55*), we likewise found that the transcriptome of mutant non-Tfh cells were concentrated in the CXCR5^+^Bcl-6^+^ gene list (Fig. S11B). Therefore, these observations suggest that *Stat1*^Mut/WT^ non-Tfh cells had a transcriptome profile more similar to that of WT Tfh cells, and also imply that gain-of-function of STAT1 endows non-Tfh cells with Tfh-associated transcriptional program.

Further, we compared the transcriptomic disparities in Tfh cells between WT and *Stat1* ^Mut/WT^ mice (Fig. 6E), and found that mutant Tfh cells highly expressed signature genes that positively regulate Tfh cells development and function, including *Ascl2*(*56*), *Maf*(*57*), *Il21*, and *Sostdc1*(*50*). On the contrary, the levels of transcripts characteristically expressed by Th2 cells (*Il4*), Th17 cells (*Il23r*, *Ccr6*, *Il17rb*, *Il17a*, *Rorc* and *Rora*), and Treg cells (*Foxp3*, *Il2a* and *Gpr83*) were decreased in mutant Tfh cells compared with controls. More importantly, Tfh cells from *Stat1* ^Mut/WT^ mice displayed high expression of master transcription factors for Th1 differentiation (*Tbx21*), as well as Th1 effector cytokines (*Ifng*, *Tnf*, *Eomes* and *Gzmk*), cytokine receptors (*Il12rb2*), and their signature chemokine receptors (*Cxcr3* and *Ccr5*). Thus, these results demonstrate that STAT1 GOF promotes the development and differentiation of Tfh cells that share phenotypic features with Th1 lineage cells.

Accumulating evidence indicates that Tfh cells with distinct states and cytokines profiles are induced based on the nature of the immune stimulus and the context within the host(*58*). Given these, we performed transcriptomic analysis on non-Tfh cells and Tfh cells from WT and *Stat1* ^Mut/WT^ mice at 20 weeks during which period the mutant mice have developed typical autoimmunity. Principal component analysis (PCA) segregated WT and mutant cells into distinct clusters, showing that both non-Tfh and Tfh cells from WT and *Stat1*^Mut/WT^ were transcriptionally different (Fig. 6F). The expression of genes involved in the IFN-I signaling pathway was increased in either non-Tfh cells or Tfh cells from mutant mice compared with that from WT mice, respectively. In addition to the increased expression of Th1 associated genes in non-Tfh cells of the mutant mice, the genes that promote the development, differentiation, and function roles of Tfh cells, such as *Cxcr5*, *Ascl2*, *Il21*, *Zeb2*, *Maf*, *Tigit*, *Icos* and *Egr2*, were also significantly upregulated in mutant non-Tfh cells compared with those of the WT group. Moreover, consistent with the above, the Th1 lineage associated genes (*Tbx21*, *Gzma, Eomes*, *Gzmk*, *Ifng*, *Il12rb2, Ccl5* and *Gzmb*) were highly expressed in Tfh cells of the mutant group, whereas the expression of Th2- (Il4), Th17-related genes (*Ccr6*, *Il17rb*) and Treg-related genes (*Ebi3*, *Il2ra*, *Lrrc32 and Foxp3*) was suppressed in the mutant Tfh cells (Fig. 6G). Collectively, these results indicate that the gain-of-function mutation of STAT1 promotes the differentiation of Tfh cells with Th1 features, which is closely similar to previously described Tfh1, Tfh1-like, or Group 1 Tfh cells(*59*).

### STAT1 T385M Directly Binds to Tfh- and Th1-Relevant Genes

To further investigate genome-wide occupancy of wild-type STAT1 and T385M mutants in the Tfh cells, we analyzed STAT1 target genes by Cleavage Under Targets and Tagmentation (CUT & Tag) coupled with high throughput sequencing. Overall, the vast majority fractions of peaks were within 3 kb upstream and downstream of the transcription start sites (TSS), with approximately half of these peaks were localized in the promoter regions of binding genes in both non-Tfh cells and Tfh cells from WT or mutant mice (Fig. 7A, Fig. S12A and table S2). Further analysis revealed the classical gamma-activated sequence (GAS)(*60*), a small palindromic sequence (5’-TTCCNGGAA-3’), as the consensus binding motif of STAT1 or T385M mutants in non-Tfh cells and Tfh cells (Fig. 7B, Fig. S12B). These data indicate that T385M gain-of-function mutation does not alter the pattern of STAT1 binding to DNA.

**Fig. 7.**
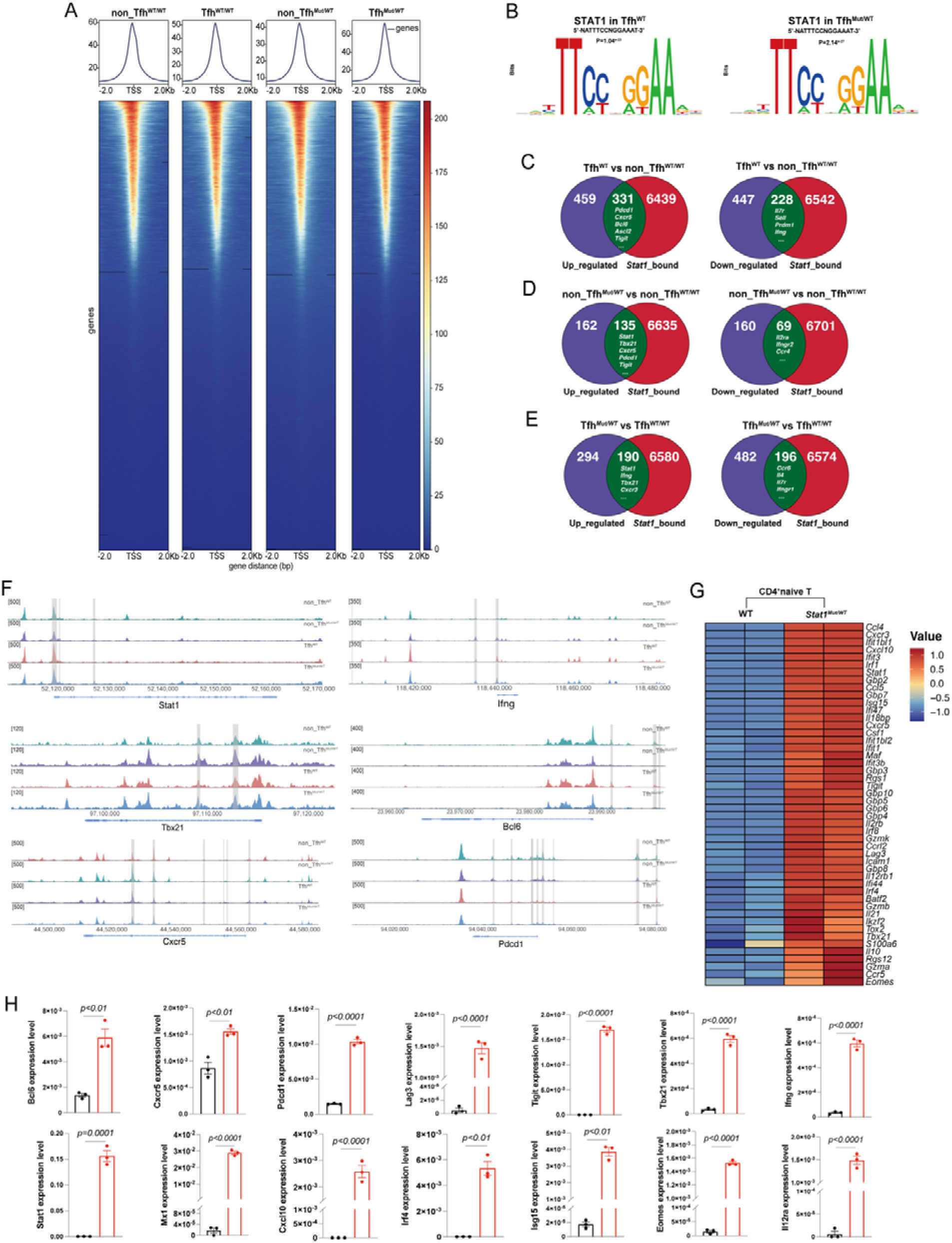
STAT1 directly binds and transcriptionally regulates Tfh- and Th1-relevant genes. (**A**) Heatmap for genome-wide distribution of STAT1 and mutant STAT1 peaks at TSS in non-Tfh ang Tfh cells. (**B**) The STAT1 and mutant STAT1 binding site is identical to the *GAS* motif. (**C-E**) Venn diagrams showing the genes regulated by STAT1 (mutant STAT1) and STAT1(mutant STAT1)-bound genes between different cells as indicated. Differentially expressed genes were marked with purple. Differentially bound genes were marked with red. (**F**) Tracks of STAT1 (mutant STAT1)-binding peaks located at gene loci including *Stat1*, *Ifng*, *Tbx21*, *Cxcr5*, *Pdcd1* and *Bcl6*. (**G**) Heatmap for DEGs of CD4^+^ naïve T cells between WT and *Stat1*^Mut/WT^ mice. (**H**) CD4^+^CD25^-^CD44^-^CD62L^+^ naïve CD4^+^T cells isolated from WT and *Stat1*^Mut/WT^ mice, and the mRNA expression level of indicated genes was determined by quantitative PCR. Data were represented as means ± SEM and analyzed with two-tailed unpaired *t* test.

Integrating STAT1-bound genes in WT Tfh cells with the differentially expressed gene between non-Tfh and Tfh cells, we found that 331 STAT1-bound genes were directly positively regulated by STAT1 in Tfh cells, including *Cxcr5*, *Bcl6*, *Pdcd1*, *Ascl2* and *Tigit* (Fig. 7C, data file S4). We next compared the STAT1-bound genes with those differentially regulated by it between WT non-Tfh and mutant non-Tfh cells, and found that the gain-of-function mutation of STAT1 also directly promoted the expression of key genes involved in Tfh differentiation and function, such as *Cxcr5*, *Pdcd1*, and *Tigit* (Fig. 7D, data file S4). More interestingly, mutant STAT1 in Tfh cells directly bound to key regulatory genes associated with the Th1 lineage, such as *Stat1*, *Tbx21*, *Cxcr3* and *Ifng*, with a significantly higher binding intensity than that in WT Tfh cells (Fig. 7E-F, data file S4). Thus, mutant STAT1 might directly bind to the *cis*-GAS element in the promoter of both Tfh and Th1 signature genes to promote the Tfh1-like cell differentiation.

Subsequently, we analyzed the effects of STAT1 on the differentiation of Th1, Th2, and Th17 cells. As shown in Figure S13A-B, compared with WT, mutant STAT1 promoted Th1 differentiation and induced *Tbx21* and *Ifng* expression. Under Th2-polarized context, mutant STAT1 had no impact on the Th2 polarization but inhibited *Il4* and *Gata3* levels (Fig. S13C-D). In agreement with previous studies, the differentiation of naïve T into Th17 were obviously impaired in *Stat1* T385M mice, accompanied by the reduction in the expression of *Il17a* and *Rorc* genes (Fig. S13E-F). In addition, mutant STAT1 slightly inhibited the induction of iTreg with decreased *Foxp3* expression (Fig. S13G-H). Consistent with recent report(*47*), hyperactive STAT1 had no effects on the suppressive activity of Treg cells *in vitro*, hinting that the subtle reduction in Treg cell differentiation may not be the main cause of autoimmunity. Therefore, STAT1 GOF promotes the polarization of Th1 and induces the Th1-relavant gene expression.

To explore whether STAT1 GOF confers naive T cells more susceptible to differentiating into Tfh1 subsets, we performed RNA-Seq on naive T cells isolated from both WT and mutant mice. In addition to the increased expression of type I interferon signaling pathway genes, the transcriptomic levels of Tfh- (*Cxcr5*, *Tigit*, *Irf4, Maf*) and Th1-relevant (*Tbx21*, *Ifng*, *Cxcr3, Gzmk*) genes which directly bound by STAT1 was significantly upregulated in naive T cells of mutant mice (Fig. 7G).

Furthermore, real-time PCR similarly confirmed the expression of key regulatory genes for Tfh and Th1 was increased in naive T cells of mutant mice compared with the control (Fig. 7H). These results suggest that STAT1 GOF may initially predispose naive T cells with transcriptomic characteristics of Tfh1 cells.

### STAT1 T385M Persistently Drives and Maintains Tfh1 Differentiation

Based on the aforementioned alterations, we detected IFN-γ production in cTfh from STAT1-GOF patients. We found that compared with healthy controls, the frequency of IFN-γ-expressing cells in cTfh from STAT1-GOF patients was significantly increased (Fig. 8A), and the intensity of T-bet cells was also markedly higher than those in healthy controls (Fig. 8B). Consistent with previous studies, the proportion of IL-17-producing cells was reduced in STAT1-GOF patients (Fig. 8A)(*44, 45, 61*). Either at 20-22 weeks (Fig. 8C) or 6-8-weeks (Fig. S14A), the proportion of IFN-γ^+^ cells in Tfh cells of mutant mice was also significantly higher than that in littermate control.

**Fig. 8.**
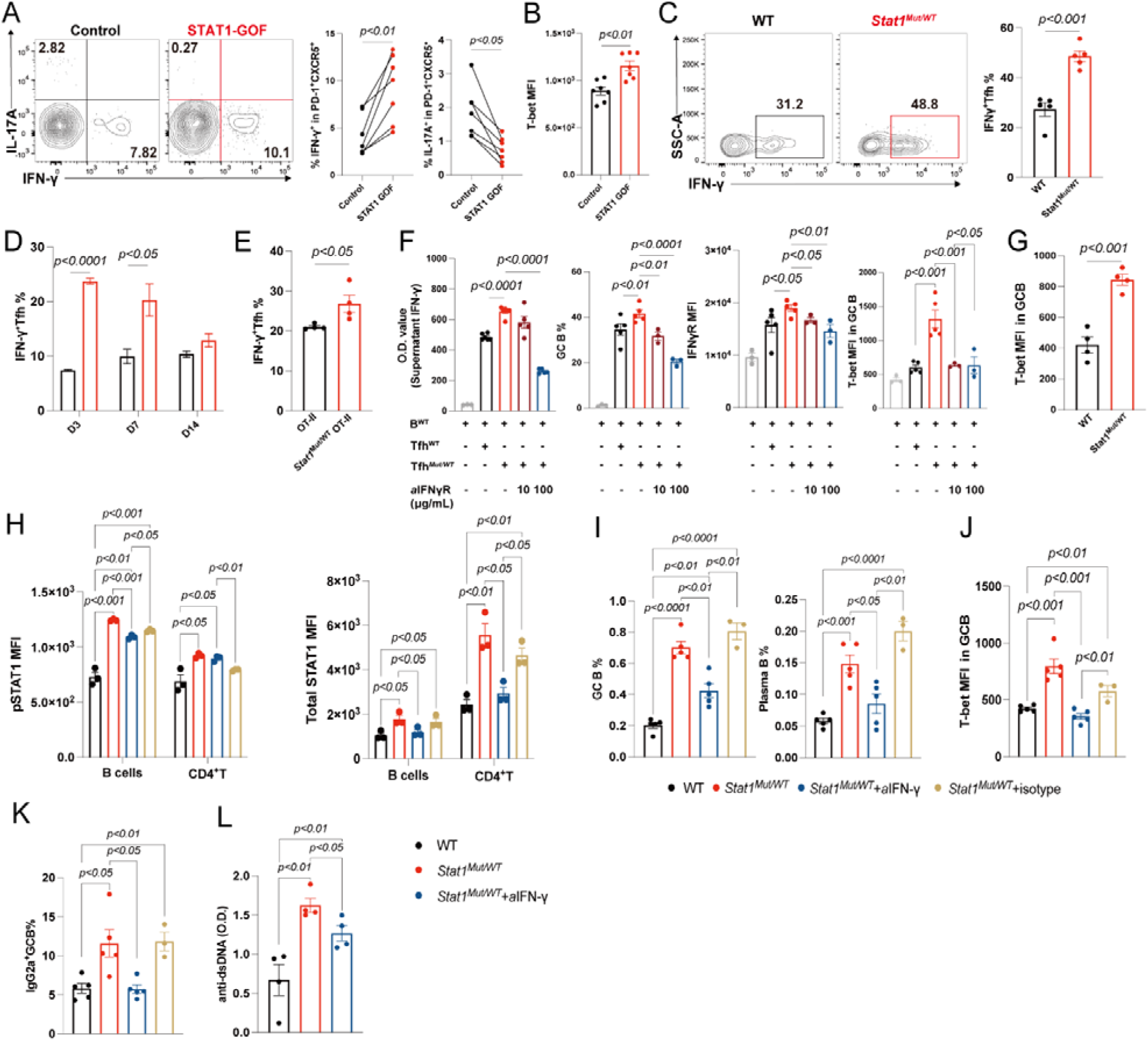
IFN-γ neutralization effectively ameliorates Tfh1 expansion and autoimmunity in STAT1 GOF mice. (**A**) Representative flow cytometry plots and pooled data of the percentage of IFN-γ^+^ and IL-17A^+^ in CD3^+^ CD4^+^CXCR5^+^PD-1^+^ cTfh cells from *STAT1*-GOF patients and controls. (**B**) MFI of T-bet in cTfh cells from *STAT1*-GOF patients and controls. (**C**) Representative flow cytometry plots and pooled data of the percentage of IFN-γ^+^ in splenic Tfh of mice at age of 20-22 weeks. (**D**) Percentages of IFN-γ^+^ in Tfh of mice after immunization with KLH/CFA for indicated time points. (**E**) Percentage of IFN-γ^+^ in Tfh from indicated donor cells. (**F**) Isolated Tfh from indicated mice and B cell from WT co-cultured *in vitro*. IFN-γ level in supernatant, percentage of GC B cells and IFNγR, T-bet MFI in GC B cells were determined. (**G**) T-bet MFI in splenic GC B cells of mice at age of 6-8 weeks. (**H**) MFI of pSTAT1 (Y701) and total STAT1 in B cells and CD4^+^T cells from mice injected with anti-IFN-γ antibody or isotype control. (**I**) Percentage of splenic GC B cells and plasma B cells in indicated mice. (**J**) MFI of T-bet in splenic GC B cells in indicated mice. (**K**) Percentage of IgG2a^+^ in GC B cells in indicated mice. (**L**) Serum anti-dsDNA antibody level in indicated mice was determined. Each symbol represents an individual throughout. Data were represented as means ± SEM and analyzed with two-tailed paired(A) or unpaired *t* test(B-L).

Previous research has shown some overlap or competition in specific Tfh and Th1 effector CD4^+^ T cell subsets(*9, 10, 53, 62–65*). The precise mechanisms controlling Tfh and Th1 cell divergence are incompletely understood. Thus, we further determine the kinetics of IFN-γ-producing Tfh cells and found that at different time points following KLH immunization, the proportion of IFN-γ-producing cells within Tfh cells of mutant mice was consistently higher than that in control mice (Fig. 8D). Moreover, on the 7th day post-immunization, the abundance of IFN-γ^+^ Tfh cells in mutant mice was even higher than that in non-Tfh cells (Fig. S14B). Moreover, the ratio of *Tbx21* to *Bcl6* in Tfh cells of mutant mice is consistently significantly higher than that in WT (Fig. S14C). In addition, in OT-II transferred model, the percentage of IFN-γ+ Tfh cells was also elevated in *Stat1*^Mut/WT^ mice (Fig. 8E). Together, these results indicate that STAT1 T385M intrinsically persistently drives IFN-γ production and maintains the Tfh1 features.

### Ruxolitinib Effectively Reverses the Autoimmune Phenotype of *Stat1*-GOF mice

IFN-γ binds to its receptor and signals through JAKs/STAT1 to initiate and establish a Th1 cell-associated gene-expression profiles in a positive feedforward manner(*66–68*). Emerging evidence suggest that JAKs inhibitors are both safe and effective in the treatment of *STAT1*-GOF patients with severe autoimmunity and immune dysregulation(*69–71*). To determine the effects of JAKs inhibitors on autoimmunity and humoral immunity in *Stat1*^Mut/WT^ mice, we administered Ruxolitinib, which inhibits JAK1 and JAK2, to mutant mice via intragastric gavage. Compared with the placebo group, after 2 weeks of administration, the levels of phosphorylated STAT1 and total STAT1 in CD4^+^T cells and B cells of the mice were significantly reduced and even reverted to WT control (Fig. S15A), indicating that Ruxolitinib can effectively inhibit STAT1 activity. In addition, Ruxolitinib-treated mice had smaller spleen size, fewer splenocytes (Fig. S15B), and lower early CD44^+^CD62^-^ effector T cells in spleen than mice treated with vehicle (Fig. S15C). Of note, at the end of treatment, Ruxolitinib resulted in a significant decrease in the proportion and number of splenic CXCR5^+^PD-1^+^ Tfh cells (Fig. S15D), CXCR5^+^Bcl-6^+^ Tfh cells (Fig. S15E), as well as the IFN-γ-producing CXCR5^+^Bcl-6^+^ Tfh1 cells (Fig. S15F), accompanied by a reduction in the frequency and absolute number of splenic GC B cells and plasma cells (Fig. S15G-H), and these indices even reverted to the levels of normal mice. Meanwhile, the level of anti-dsDNA IgG in serum in mutant mice was also restored to the normal level (Fig. S15I). However, serum IgG2a and IgG2c in Ruxolitinib-treated mice remained a comparable level to *Stat1*^Mut/WT^ mice (Fig. S15J). Thus, these above data suggest that inhibiting JAKs activity *in vivo* can almost completely reverse the autoimmunity caused by STAT1 GOF.

### IFN-**γ** Neutralization Alleviates Autoimmunity of *Stat1*-GOF mice

Based on results above, we hypothesized that the excessively produced IFN-γ by Tfh1 might be the underlying mechanism for disease pathogenicity and spontaneous autoimmunity in *Stat1*^Mut/WT^ mice. Accordingly, Tfh cells isolated from WT or mutant mice individually, and B cells were cocultured in the absence or presence of IFNγR-neutralizing antibody(αIFN-γ) *in vitro*. After 6 days of culture, compared with Tfh cells from WT mice, Tfh cells from the mutant mice produced a higher level of IFN-γ in the culture supernatant, induced the formation of GC B at a higher proportion, while blocking the IFNγR with αIFNγR suppressed the increased GC formation in dose-dependent manner (Fig. 8F). Furthermore, it is worth noting that the mRNA expressions of both *Ifngr1* and *Ifngr2* on the Tfh cells from the mutant group were significantly downregulated (Fig. 6E). However, we found that mutant Tfh cells promoted the IFNγR expression in GC B cells (Fig. 8F). Consistently, GC B cells induced by mutant Tfh cells *in vitro* exhibited higher expression of T-bet (Fig. 8F). Likewise, the level of the T-bet in GC B cells from mutant mice was also higher than that in control in basal state (Fig. 8G). There results suggest that IFN-γ secreted by Tfh cells from the mutant group may act on GC B cells with a paracrine manner.

To further demonstrate whether the massive production of IFN-γ causes autoimmunity in mutant mice, 6-week-old *Stat1*^Mut/WT^ mice, which had already developed anti-dsDNA antibody, were intraperitoneally administrated with αIFN-γ or isotype control antibodies. Following 3 weeks consecutive treatment, αIFN-γ blockade led to significant reduced levels of pSTAT1 and total STAT1, along with a smaller spleen size and lower absolute number of splenocytes (Fig. 8H, Fig. S16A-B). αIFN-γ treated mice had reduced frequency and numbers of CD44^+^ effector T cells (Fig. S16C), plasma cells, GC B cells and lower T-bet expression in GC B cells (Fig. 8I) than in mice treated with isotype controls. In agreement with the roles for IFN-γ in switching to IgG2a/c in murine, percentage of IgG2a^+^ GC B cells and serum IgG2c level were also reduced in αIFN-γ neutralization mice (Fig. 8J-K). In addition, anti-dsDNA antibody levels in αIFN-γ-treated mutant mice were significantly declined comparable with the concentration found in *Stat1*^Mut/WT^ mice (Fig. 8L). Thus, these results indicate that overproduced IFN-γ derived from Tfh1 leads to autoimmunity and defective humoral immune responses, and neutralizing IFN-γ cytokines display an effective strategy for autoimmunity in STAT1 GOF model.

## DISCUSSION

Inborn errors of immunity (IEI) always shed light on genes and signaling pathways critical for human immunity. Since its initially discovered in 2011(*44, 45*), about half of the STAT1 GOF patients exhibit various autoimmune manifestations, such as hypothyroidism, enterocolitis, immune cytopenia, autoimmune hepatitis, endocrinopathies, and systemic lupus erythematosus, but the underlying pathogenic mechanism remains to be elusive. In this study, with a cohort of *STAT1*-GOF patients from multiple regions of China and a corresponding mutant mouse model (*Stat1* T385M), we analyzed how gain-of-function mutation in STAT1 drives autoimmunity and discovered that overactivated STAT1 can directly target Tfh and Th1 cell signature genes and thereby promote the development of Tfh1 cells with excessive IFN-γ production, which implicated in autoantibody production and the development of autoimmunity.

It has long been recognized that the IFN-γ-dependent activation of STAT1 is essential for inducing the expression of T-bet to establish Th1 cell-associated gene program and to guide commitment toward Th1 fate(*66–68*). Indeed, patients with loss-of-function mutation in STAT1 show more susceptibility to intra-macrophagic mycobacteria, BCG vaccine, and viruses partly due to impaired Th1 differentiation and IFN-γ signaling(*72, 73*). Likewise, our data revealed that overactivated STAT1 promoted naïve CD4^+^ T cell differentiation into Th1 cells with increased T-bet expression and IFN-γ production in Th1 polarizing condition *in vitro*. Unexpectedly, we also found that either in different foreign immunization or in endogenous immune context, hyperactive STAT1 promoted the differentiation of Tfh1 cells, which is characterized by a typical Tfh cells features, such as highly expressing CXCR5, PD-1, Bcl-6, IL-21 and localizing within follicles. Meanwhile, these Tfh cells also exhibit Th1 cell characteristics, such as high expression of T-bet, IFN-γ, CXCR3 and other related molecules. Previous work has suggested that Tfh and Th1 share a transitional stage, and IFNγ-producing Tfh and Th1 cells develop simultaneously in mice mediated by IL-12-STAT4(*40, 62, 63, 65*). However, these cell features were not sustained during later phase of Th1 cell differentiation because of enforced T-bet antagonize expression of Bcl-6 to low levels and eventually allows for Th1 cell specification at the expense of the Tfh-Th1 like state(*74–76*). Emerging evidence suggests that environmental cytokines determine the Tfh versus Th1 cell fates and regulate flexibility in these cells by modulating the balance of the T-bet and Bcl-6(*74–76*). Importantly, it is reported that STAT1 is required for IL-6–mediated Bcl-6 induction for early but not necessary for the complete differentiation of Tfh cells(*42*). Accordingly, we speculated that overactivated STAT1 may perceive and integrate diverse environmental signals, such as IFN-γ, IL-12, IFN-α/β, IL-6, or IL-21, to synchronously induce the expression of T-bet and Bcl-6 to tip the balance between their expression, thereby favoring cells to differentiate toward Tfh1 and sustaining their hybrid characteristics. Indeed, our data suggested that IFN-γ expression was persistently elevated in Tfh cells from mutant mice, and the T-bet/Bcl-6 ratio was also significantly higher compared with the control group.

IFN-γ^+^ Tfh cells have been detected in various scenarios, including viral infection (e.g., LCMV, influenza, *ZIKV*), Malaria or *Toxoplasma gondii* challenging, vaccination with inactivated viral vaccines(*77–80*). In addition, the proportion of IFN-γ^+^ Tfh cells is increased in certain autoimmune diseases (e.g., SLE) and common variable immunodeficiency (CVID)(*37, 81, 82*). However, the role of IFN-γ^+^ Tfh cells has long been controversial. For example, severe malaria infection induces the development of a population of CXCR5^int^PD-1^int^ pre-Tfh cells that share phenotypic features with Th1 lineage cells, which compromised the differentiation of mature Tfh cells and the GC reaction(*77*). Rather, it is reported that T-bet^+^ Tfh cells is required for Tfh function and also critical for the proper maturation of GC B cells appropriate to viral challenge(*40, 83*). Similarly, another study has shown that after influenza infection, at the peak of the response Tfh cells transiently produce IFN-γ, which is indispensable for the differentiation of influenza-specific lung-resident memory B cells responses against heterosubtypic infection(*80*). In the present study, our data demonstrated that hyperactive STAT1 lead to the sustained excessive production of IFN-γ^+^ Tfh cells, which not only compromised humoral response specially in antibody class switching but also facilitated the progression of pathological autoimmunity. Accordingly, we supposed that the optimal and transient formation of IFN-γ□ Tfh cells may be crucial for the host immune response against viral or another foreign antigen. However, their persistent and excessive production is likely to induce pathological immune responses, such as autoimmunity and multi-organ inflammatory damage(*31*). Of course, the characteristics of IFN-γ□ Tfh cells, the differences between IFN-γ□ Tfh and IFN-γ□ Tfh cells, as well as the similarities and discrepancies of IFN-γ□ Tfh cells between normal mice and STAT1 GOF mice remain undefined, and further work using reporter mice or fate-mapping approaches are therefore required to elucidate.

Utilizing another STAT1 GOF mouse model (R274Q), a recent study has shown that deletion of IFNγR, rather than IFNαR, reverses autoimmunity induced by STAT1 GOF(*47*). Consistent with this finding, our present study revealed that neutralization of IFN-γ abrogated STAT1-driven excessive Tfh1 cell frequencies, autoantibody production and spontaneous autoimmunity. A question that emerges is how uncontrolled IFN-γ from Tfh1 cells induce autoimmunity. In the GC microanatomical niche, IFN-γ from Tfh may drive autoimmunity by acting on Tfh cells themselves or GC B cells via autocrine or paracrine mechanisms, respectively. Our *in vitro* co-culture assay results showed that IFNγR blockade attenuated mutant Tfh cells-induced GC formation. In addition, blocking IFN-γ signaling *in vivo* robustly abrogated the elevated T-bet levels in GC B cells of mutant mice. Further, both *Ifngr1* and *Ifngr2* expressions were significantly downregulated in mutant Tfh cells either in immune or in basal state. Seminal studies have illustrated that B cell–intrinsic IFNγR/T-bet signaling is central to spontaneous autoimmune GC formation and autoimmunity(*84, 85*).

Collectively, these implied that IFN-γ from Tfh1 cells may primarily act on GC B cells to induce T-bet in GC B via IFNγR, and that the interplay between GC B -Tfh1 cells promotes autoimmunity and IgG2-skewed humoral response. However, the precise mechanism by which the IFN-γ-T-bet or IFN-γ-STAT1-T-bet axes regulated autoimmunity need to be further better clarified.

*STAT1*-GOF patients show enhanced STAT1 phosphorylation in response to IFN I/II due at least in part to impaired or delayed dephosphorylation and increased expression of total STAT1(*86–88*). In terms of autoimmune or immune dysregulated manifestations, *STAT1*-GOF patients were commonly treated with immunosuppressive agents, which have severe side effects. Hematopoietic stem cell transplantation (HSCT) is a curative treatment to correct the underlying genetic defect in STAT1-GOF patients(*89*). However, HSCT increases the risk of secondary graft failure, and approximately 40% of patients die from secondary graft failure or other complications following HSCT(*72, 73, 89*). With the advent of JAK inhibitors (JAKi), such as Ruxolitinib and Baricitinib (both primarily inhibiting JAK1/2 kinases), Ruxolitinib is considered to alleviate the enhanced IFN I/II signaling and therefore the autoimmunity caused by *STAT1* GOF mutations(*70, 71*). Furthermore, Ruxolitinib can be employed as a bridging therapy for *STAT1*-GOF patients awaiting HSCT, with the aim of reducing post-transplant complications(*90*). However, during treatment with JAKi, the risk of infection in patients may increase. Our research indicated that anti-IFN-γ monoclonal antibodies (e.g., Emapalumab, Fontolizumab) may provide a novel approach for the treatment of autoimmunity in patients with *STAT1* GOF mutations, particularly for more severe cases of autoimmunity.

In summary, our data show that gain-of-function mutation in *STAT1* intrinsically promotes Tfh1 differentiation, which contributes to autoantibody production,spontaneous autoimmunity and defective humoral immunity. Our findings underscore the importance of STAT1 in humoral immunity mediated by Tfh and GC B cells. Our results further imply that STAT1 activity should be finely tuned within a reasonable magnitude to facilitate optimal immune responses while prevents excessive responses to tonic stimuli which induce the occurrence of autoimmunity.

## Supporting information

Graphical abstract

## Acknowledgement

We thank all the patients, their families, and healthy volunteers for their participation in the study. We thank all the members of Chongqing Key Laboratory of Child Rare Diseases in Infection & Immunity for discussion.

## Data and materials availability

Data generated in this study have been deposited in the OMIX database. The dataset comprising CUT&Tag sequencing data of genome-wide STAT1 chromatin binding in sorted splenic Tfh and non-Tfh CD4□ T cell subsets was under accession number OMIX013580. The dataset including gene expression matrices for sorted splenic Tfh and non-Tfh CD4□ T cell subsets of mice aged at 20 weeks or immunized with KLH/CFA was under accession number OMIX013582. Expression data from BCL6-YFP-positive Tfh cells, BCL6-YFP-negative Tfh cells, and non-Tfh cells were downloaded from GSE106138. Tabulated data underlying the figures are provided in data file S5. All data needed to evaluate the conclusions in the paper are present in the paper or the Supplementary Materials.

## Author contribution

**Ran Chen**: Data curation, Formal analysis, Investigation, Methodology, Software, Validation, Visualization, Writing – original draft, Writing – review & editing. **Yanjun Jia**: Conceptualization, Data curation, Funding acquisition, Investigation, Methodology, Project administration, Resources, Supervision, Validation, Writing – original draft, Writing – review & editing. **Yunfei An**:

Conceptualization, Project administration, Resources, Supervision, Validation, Writing – review & editing. **Xiaodong Zhao**: Conceptualization, Project administration, Resources, Supervision, Validation, Writing – review & editing. **Xuemei Chen**: Data curation, Investigation. **Jigui Yang**: Data curation. **Huilin Mu**: Investigation. **Shengqiao Mao**: Investigation. **Siyu Chen**: Investigation. **Rui Gan**: Investigation. **Qinglv Wei**: Investigation. **Wenjing Tang**: Investigation. **Junfeng Wu**: Investigation. **Wuyang He**: Investigation. **Satoshi Okada**: Writing – review & editing. **Lina Zhou**: Investigation.

## Supplementary Materials

**Fig. S1.** Identification of two novel STAT1-GOF mutants

**Fig. S2.** Enhanced STAT1 signaling in STAT1 GOF mice

**Fig. S3.** T cells development in thymus of STAT1 GOF mice

**Fig. S4.** Expanded and activated CD4+T cells in STAT1 GOF mice at age of 6-8 weeks

**Fig. S5.** Expanded and activated CD4+T cells in STAT1 GOF mice at age of 20-22 weeks

**Fig. S6.** Expanded Tfh cells in lymph nodes of STAT1 GOF mice

**Fig. S7.** Tfh from STAT1 GOF mice induced autoantibody production

**Fig. S8.** Expanded Tfh cells in STAT1 GOF mice infected with LCMV Armstrong

**Fig. S9.** Intrinsic expansion of Tfh cells from STAT1 GOF mice

**Fig. S10.** Defective humoral responses in different immune challenge, related to Figure 5

**Fig. S11.** Cell sorting strategy and GSEA analysis, related to Figure 6

**Fig. S12.** Distribution of STAT1 binding profile in gene loci

**Fig. S13.** Dysregulated CD4+T cells differentiation in vitro

**Fig. S14.** Tfh1 skewing phenotype in STAT1 GOF mice, related to Figure 8.

**Fig. S15.** Autoimmunity phenotype in STAT1 GOF mice reverted by Ruxolitinib treatment

**Fig. S16.** Alleviated STAT1 hyperactivity and CD4+T cells activation following αIFN-γ treatment, related to Figure 8.

**Table S1.** Clinical and immunological manifestations of STAT1-GOF patients Table S2. Summary of STAT1 Binding Peak Positions

Data file S1. Supplementary materials

Data file S2. Primer sequencesd

Data file S3. Different genes expression between non-Tfh and Tfh

Data file S4. Gene lists of RNA-Seq and CUT&Tag combined analysis

Data file S5. Tabulated data

