## Supplementary material for "Hyperactive STAT1 Promotes T Follicular Helper Type 1 Cell Differentiation to Trigger Autoimmunity": Data file S1-S5: Data file S1 Supplementary materials.docx

| Reagent or resource | | Source | | Cat# | Clone | |
| --- | --- | --- | --- | --- | --- | --- |
| **Antibodies** | |  | |  |  | |
| Anti-human CXCR5 BV421 | | Biolegend | | 356920 | J252D4 | |
| Anti-human PD-1 FITC | | Biolegend | | 329904 | EH12.2H7 | |
| Anti-human CXCR3 APC | | BD bioscience | | 561732 | 1C6 | |
| Anti-human CCR6 PE | | Biolegend | | 353410 | G034E3 | |
| Anti-human CD4 PE-cy7 | | Biolegend | | 300512 | RPA-T4 | |
| Anti-human CD45RA FITC | | BD bioscience | | 555488 | HI100 | |
| Anti-human IFN-γ APC | | Biolegend | | 502512 | 4S.B3 | |
| Anti-human CD3 Percp | | Biolegend | | 300326 | HIT3a | |
| Anti-Total Stat1 (N-Terminus) PE | | BD bioscience | | 558537 | 1/Stat1 | |
| Anti-Stat1 (pY701) PE | | BD bioscience | | 612564 | 4a | |
| Anti-mouse CD3e BV510 | | BD bioscience | | 563024 | 145-2C11 | |
| Anti-mouse CD8a PE-cy7 | | BD bioscience | | 552877 | 53-6.7 | |
| Anti-mouse CD4 FITC | | Biolegend | | 100406 | GK1.5 | |
| Anti-mouse CD4 APC | | BD bioscience | | 553051 | GK1.5 | |
| Anti-mouse CXCR5-biotin | | BD bioscience | | 551960 | 2G8 | |
| APC-streptavidin | | Jackson Immuno | | 016-130-084 | —— | |
| Anti-mouse CD44 BV510 | | Biolegend | | 103044 | IM7 | |
| Anti-mouse CD44 APC | | Biolegend | | 103012 | IM7 | |
| Anti-mouse CD62L FITC | | BD bioscience | | 561917 | MEL-14 | |
| Anti-mouse CD69 BV421 | | Biolegend | | 104545 | H12F3 | |
| Anti-mouse CD25 BV421 | | Biolegend | | 102034 | PC61 | |
| Anti-mouse PD-1 (CD279) PE-cy7 | | Biolegend | | 109109 | RMP1-30 | |
| Anti-mouse Foxp3 PE-cy5 | | Invitrogen | | 15-5773-82 | FJK-16s | |
| Anti-mouse Bcl-6 PE | | BD bioscience | | 561522 | K112-91 | |
| Anti-mouse T-bet PE-cy7 | | invitrogen | | 25-5825-82 | eBio4810 | |
| Anti-mouse IFN-γ PE-cy7 | | Biolegend | | 505826 | XMG1.2 | |
| Anti-mouse IL-17A PE | | Biolegend | | 506904 | TC11-18H10.1 | |
| Anti-mouse IL-4 PE | BD bioscience | | 554435 | | | 11B11 |
| Anti-mouse IL-4 PE-cy7 | | BD bioscience | | 560699 | 11B11 | |
| Anti-mouse RORγt BV421 | | BD bioscience | | 562894 | Q31-378 | |
| Anti-mouse GATA3 BV421 | | Biolegend | | 653814 | 16E10A23 | |
| Anti-mouse B220 FITC | | BD bioscience | | 553087 | RA3-6B2 | |
| Anti-mouse GL7 Pacific Blue | | Biolegend | | 104545 | GL7 | |
| Anti-mouse CD95 PE-cy7 | | BD bioscience | | 557653 | Jo2 | |
| Anti-mouse CD138 APC | | Biolegend | | 142505 | 281-2 | |
| Anti-mouse IgG1 BV510 | | Biolegend | | 406621 | RMG1-1 | |
| Anti-mouse IgG2a PE | | Biolegend | | 407108 | RMG2a-62 | |
| Anti-mouse CD45.1 FITC | | BD bioscience | | 561871 | A20 | |
| Anti-mouse CD45.2 BV510 | | BD bioscience | | 740131 | 104 | |
| Fixable viability dye efluor780 | | Invitrogen | | 65-0865-14 | —— | |
| Anti-mouse IgD FITC | | Biolegend | | 405703 | 11-26c.2a | |
| Anti-mouse CD4 APC | | BD bioscience | | 553051 | RM4-5 | |
| Anti-mouse CD45.2 PE | | BD bioscience | | 560695 | 104 | |
| Phospho-Stat1(Tyr701) Rabbit mAb | | Cell Signaling Technology | | 7649 | D4A7 | |
| HRP-conjugated β-actin monoclonal antibody | | Proteintech | | HRP-66009 | 2D4H5 | |
| Anti-rabbit IgG, HRP-linked antibody | | Cell Signaling Technology | | 7074 | —— | |

| Reagent or resource | Source | Cat# |
| --- | --- | --- |
| **Chemicals, Peptides, Recombinant Proteins, Culture medium and Consumables** | | |
| PMA | Sigma-Aldrich | P1585 |
| Ionomycin | Sigma-Aldrich | 407952 |
| Recombinant mouse IFN-γ | PeproTech | 315-05 |
| Recombinant mouse IFN-α | RD | 10149-IF-010 |
| ProLong™ Glass Antifade Mountant | Invitrogen | P36980 |
| Recombinant mouse IL-2 | PeproTech | 200-02 |
| Recombinant mouse IL-4 | PeproTech | 214-14 |
| Recombinant mouse IL-12 p70 | PeproTech | 210-12 |
| Recombinant mouse IL-6 | PeproTech | 216-16 |
| Recombinant mouse IL-23 | RD | 1887-ML |
| Recombinant mouse IL-1β | PeproTech | 211-11B |
| Recombinant mouse TGF-β | RD | 7666-MB |
| Anti-mouse IFN-γ | RD | MAB485-SP |
| Anti-mouse IL-4 | RD | MAB404-100 |
| Recombinant mouse IL-7 | PeproTech | 217-17 |
| SBA Clonotyping System-C57BL/6-HRP | Southern Biotech | 5300-05B |
| OPD | Sigma-Aldrich | P9187 |
| CellTrace™ CFSE | Invitrogen | C34554 |
| M.Tuberculosis Des.H37 Ra | Difco | 231141 |
| Freund’s Adjuvant, incomplete | Sigma-Aldrich | F5506 |
| Anti-mouse CD3 | InVivoMAb | BE0001 |
| Anti-mouse CD28 | InVivoMAb | BE0015 |
| Anti-mouse IFN-γR | InVivoMAb | BE0029 |
| Goat anti-mouse IgM F(ab)_2_ Fragment | Jackson Immuno | 115-006-020 |
| NP(4) | LGC Biosearch Technologies | N-5050XL |
| Anti-Mouse IFNγ(12G1.N) | InVivoMAb | BE0055 |
| rat IgG1 isotype control, anti-horseradish peroxidase | InVivoMAb | BE0088 |
| NP(30) | LGC Biosearch Technologies | N-5050H |
| TRIzol™ | Invitrogen | 15596026CN |
| Glycogen, RNA grade | Thermo Scientific | R0551 |
| KLH | LGC Biosearch Technologies | K-1000-5 |
| NP-KLH | LGC Biosearch Technologies | N-5060 |
| OVA | Sigma-Aldrich | A5503 |
| GlutaMAX™ | Gibco | 35050061 |
| MEM NEAA | Gibco | 11140050 |
| 2-Mercaptoethanol | Gibco | 21985023 |
| Sodium Pyruvate | Gibco | 11360070 |
| RPMI | Gibco | C11875500BT |
| Tween-20 | DING GUO | DH358-3 |
| O.C.T. Compound | Sakura | 4583 |
| RIPA | Beyotime | P0013B |
| Phenylmethylsulfonylfluoride (PMSF) | Sigma-Aldrich | ST506 |
| Protease inhibitor cocktail | Sigma-Aldrich | P8340 |
| Nunc MicroWell 96-well plate | Thermo Scientific | 167008 |
| PVDF Membrane | Millipore | IPVH00010 |
| Ruxolitinib | MCE | HY-50856 |
| PEG300 | MCE | HY-Y0873 |
| GolgiStop Protein Transport Inhibitor | BD bioscience | 554724 |
| GolgiPlug Protein Transport Inhibitor | BD bioscience | 555029 |

| Reagent or resource | Source | Cat# |
| --- | --- | --- |
| **Critical Commercial Assay Kits** | | |
| BD Cytofix/ Cytoperm kit | BD Biosciences | 554722 |
| Foxp3/Transcription Factor Staining Buffer Set | eBioscience | 00-5253-00 |
| Phos-flow Fix Buffer I | BD Biosciences | 557870 |
| BD Phosflow Perm Buffer III | BD Biosciences | 558050 |
| TB Green^®^ Premix Ex Taq™ kit | Takara Bio | RR420W |
| Naive CD4^+^ T Cell Isolation Kit, mouse | Miltenyi Biotec | 130-104-453 |
| CD4 (L3T4) MicroBeads, mouse | Miltenyi Biotec | 130-117-043 |
| dsDNA – IIFT | EUROIMMUN | FA1572 |
| Mouse Anti-dsDNA IgG ELISA Kit | ALPHA DIAGNOSTIC | 5120 |
| Hyperactive Universal CUT&Tag Assay Kit for Illumina | Vazyme | TD903 |
| Mouse IFN-γ ELISA kit | 4A Biotech | CME0003-096 |
| PrimeScript RT Master Mix | Takara | RR036A |
