## Supplementary material for "Hyperactive STAT1 Promotes T Follicular Helper Type 1 Cell Differentiation to Trigger Autoimmunity": Data file S1-S5: Data file S2 Primer sequences.docx

| **Gene** | **Species** | **Primer type** | **Sequence** |
| --- | --- | --- | --- |
| Isg15 | Mouse | F-primer | AGCGAGCCTCTGAGCATCCTG |
|  |  | R-primer | GCGTGTCTACAGTCTGCGTCAG |
| Irf4 | Mouse | F-primer | CGATCGGCCCAACAAGCTA |
|  |  | R-primer | GCAAACACTTGCAGCTCTGAT |
| Cxcl10 | Mouse | F-primer | ACGTGTTGAGATCATTGCCAC |
|  |  | R-primer | TTCACTCCAGTTAAGGAGCCC |
| Il2ra | Mouse | F-primer | TGGCAACACAGATGGAGGAA |
|  |  | R-primer | CGTTAGGTGAATGCTTGGCG |
| Lag3 | Mouse | F-primer | CAACATCAACCAGACAGTGGC |
|  |  | R-primer | GTTGTCTAGGCGAGGGCAT |
| Eomes | Mouse | F-primer | ACAGTTCATCGCTGTGACGG |
|  |  | R-primer | CAGGGACAATCTGATGGGATGAAT |
| Stat1 | Mouse | F-primer | CCTGCTGTGCCTCTGGAATGATG |
|  |  | R-primer | TCCCTGGCTGCTGGTCCTTG |
| Mx1 | Mouse | F-primer | AAGGTCTTGGATGTGATGCGGAAC |
|  |  | R-primer | GCTGCTCTTGGATGTCCTGCTG |
| Tigit | Mouse | F-primer | GGTCCAAGAAAGCTCAGTGGC |
|  |  | R-primer | AGCAAATGAGTCCCAGCACAG |
| Cxcr5 | Mouse | F-primer | TAGGCACCAGCACAAACCTTC |
|  |  | R-primer | GGCCAGTTCCTTGTACAGGTCAT |
| Tbx21 | Mouse | F-primer | AACCGCTTATATGTCCACCCA |
|  |  | R-primer | CTTGTTGTTGGTGAGCTTTAGC |
| Ifng | Mouse | F-primer | ATGAACGCTACACACTGCATC |
|  |  | R-primer | CCATCCTTTTGCCAGTTCCTC |
| Pdcd1 | Mouse | F-primer | ACCCTGGTCATTCACTTGGG |
|  |  | R-primer | CATTTGCTCCCTCTGACACTG |
| Bcl6 | Mouse | F-primer | CCGGCACGCTAGTGATGTT |
|  |  | R-primer | TGTCTTATGGGCTCTAAACTGCT |
| Irf1 | Mouse | F-primer | CCCATCAGGAGGTTTCCTCG |
|  |  | R-primer | TAGGCACCAGCACAAACCTTC |
| β-actin | Mouse | F-primer | GTCATCCATGGCGAACTGGT |
|  |  | R-primer | TTCCTTGAACGACGACGACTTTGG |
