## Supplementary figure 1-16 for "Hyperactive STAT1 Promotes T Follicular Helper Type 1 Cell Differentiation to Trigger Autoimmunity": Supplementary figure 1-16.pdf

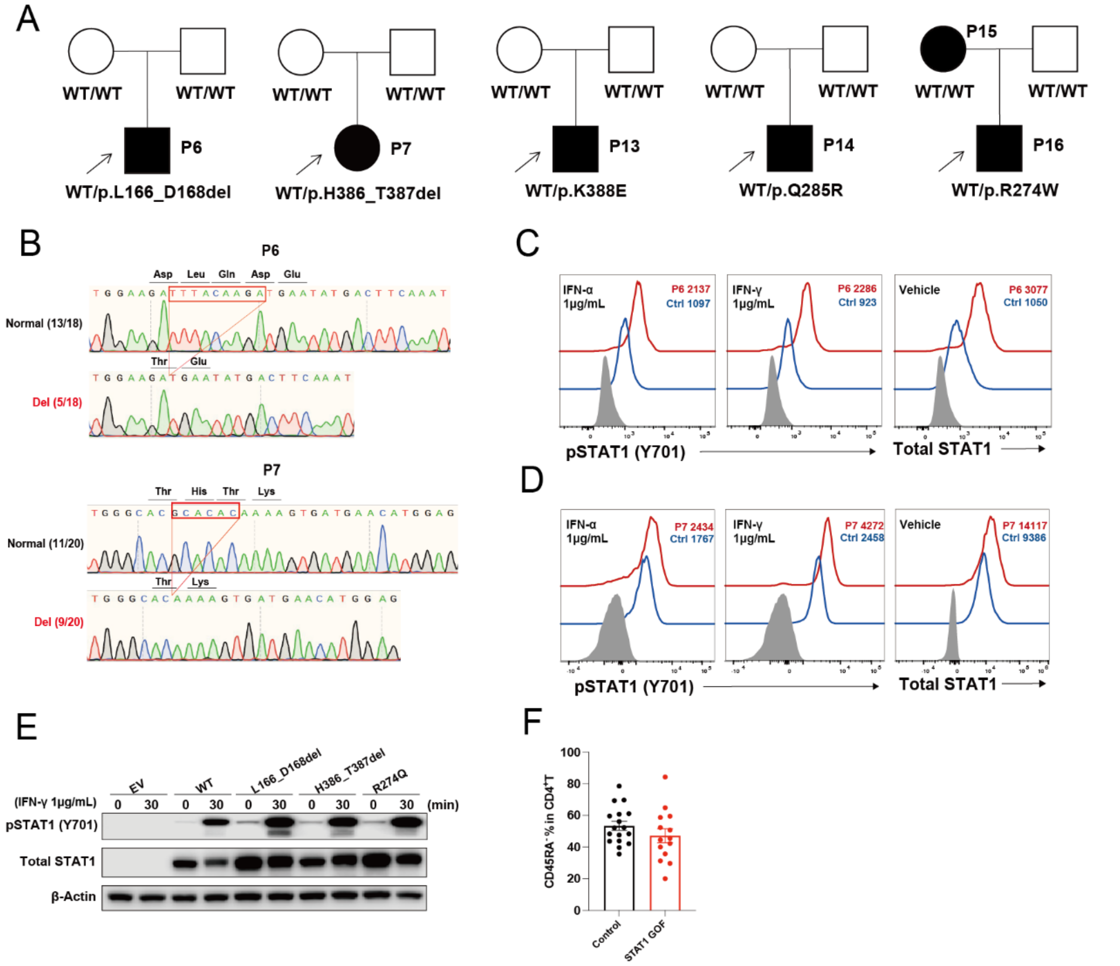


**Fig. S1 Identification of two novel *STAT1*-GOF mutants.** (**A**) *STAT1* mutations in patients of five kindreds unreported previously. (**B**) Sanger sequencing of two patients (P6 and P7) with novel STAT1 mutants. (**C-D**) pSTAT1 (Y701) induced by IFN-α (left) and IFN-γ (middle), and total STAT1 at basal level (right) of P6 (C) and P7 (D) compared to controls, respectively. (**E**) pSTAT1 and total STAT1 expression of two novel mutants in U3C cell line confirmed by western blot. (**F**) Percentage of $CD45RA^-$ in $CD4^+$T cells of *STAT1*-GOF patients and age-matched controls. Each symbol represents an individual throughout.

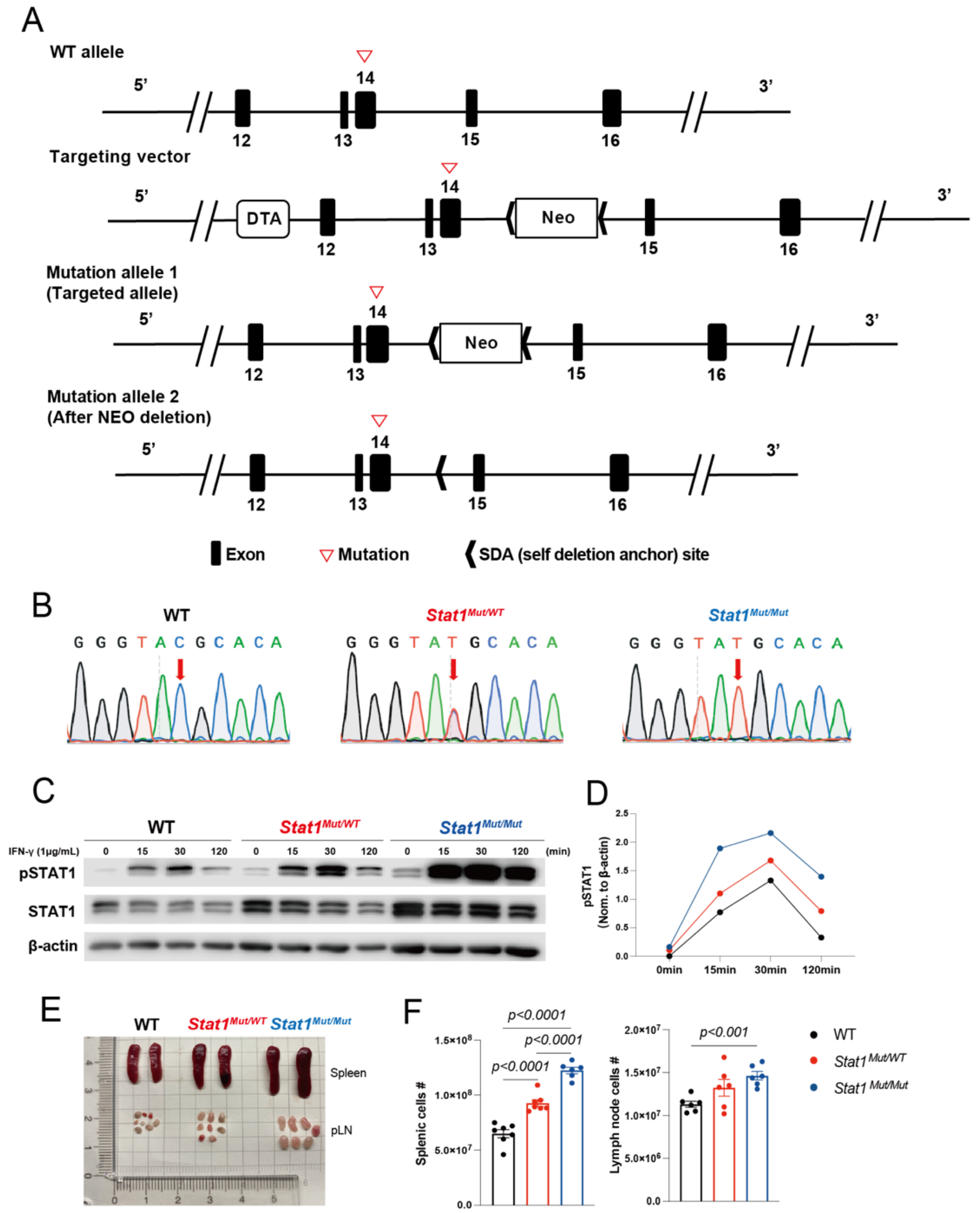


**Fig. S2 Enhanced STAT1 signaling in *Stat1*-GOF mice.**

(**A**) Schematic diagram of *Stat1* mutant mice. The targeting vector was designed to introduce the T385M point mutation (c.1154C>T) into exon 14 of the murine *Stat1* gene, utilizing 5' and 3' homology arms. A Neo cassette, flanked by SDA sites, was inserted for positive selection and subsequent removal. The construct included a DTA gene for negative selection in embryonic stem cells (ES cells). Targeted ES cell clones (C57BL/6N) were used to generate the mutant mouse model. (**B**) Sanger sequencing of WT, *Stat1*$^{\mathrm{Mut/WT}}$ and *Stat1*$^{\mathrm{Mut/Mut}}$ mice. Red arrows indicate the mutation sites. (**C-D**) pSTAT1 (Y701) and total STAT1 protein from splenic cells of mice as indicated were determined by western blot (C) and densitometric quantification (D) normalized to β-actin. (**E**) Image of enlarged spleen and lymph nodes of mice as indicated. (**F**) Numbers of splenocytes and lymph node cells of mice as indicated. Each symbol represents an individual throughout. Data were represented as means ± SEM and analyzed with two-tailed unpaired *t* test.

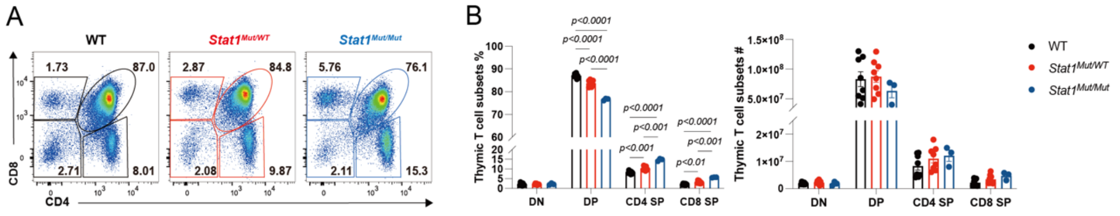

**Fig. S3 T cells development in thymus of STAT1 GOF mice.** (**A**) Flow cytometry plot of CD8 versus CD4 in thymic cells of WT, *Stat1*$^{\mathrm{Mut/WT}}$ and *Stat1*$^{\mathrm{Mut/Mut}}$ mice. (**B**) Percentages (left) and numbers (right) of DN (double negative), DP (double positive), CD4 SP (CD4 single positive) and CD8 SP (CD8 single positive) T cells of mice as indicated. Each symbol represents an individual throughout. Data are representative of three independent experiments. Data were represented as means ± SEM and analyzed with two-tailed unpaired *t* test.

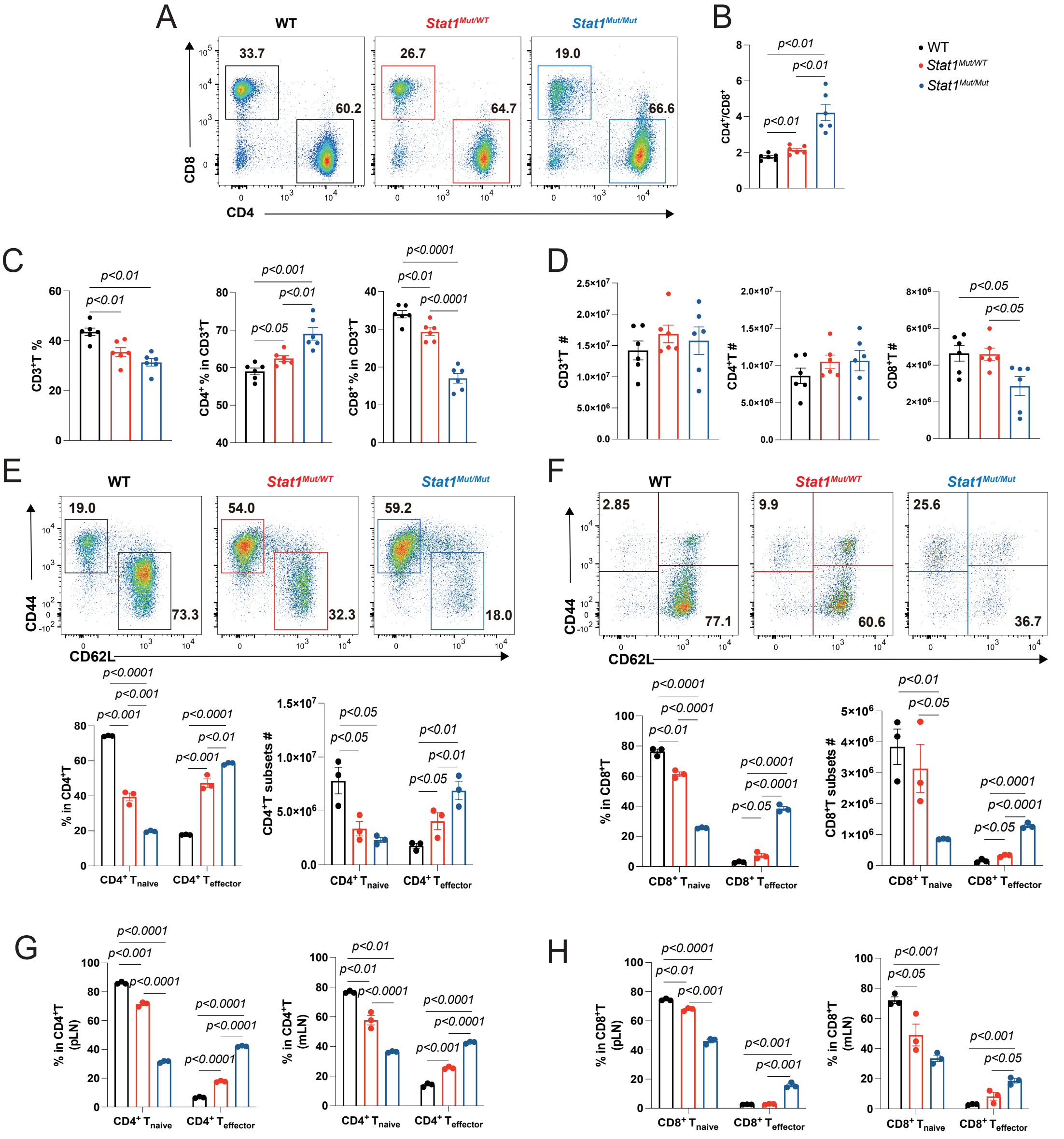

**Fig. S4 Expanded and activated CD4$^+$T cells in STAT1 GOF mice at age of 6-8 weeks.** (**A**) Flow cytometry plot of CD4 versus CD8 in splenic CD3$^+$T cells. (**B**) Ratio of CD4$^+$T to CD8$^+$T percentage. (**C**) Percentage of CD3$^+$ in live cells (left), CD4$^+$ in CD3$^+$T (middle), CD8$^+$ in CD3$^+$T cells (right) in spleen. (**D**) Numbers of splenic CD3$^+$, CD4$^+$ and CD8$^+$ T cells. (**E**) Flow cytometry plot, percentage and numbers of splenic naïve CD4$^+$T and effector CD4$^+$T cells. (**F**) Flow cytometry plot, percentage and numbers of splenic naïve CD8$^+$T and effector CD8$^+$T cells. (**G**) Percentage of CD4$^+$T subsets in pLNs and mLNs. (**H**) Percentage of CD8$^+$T subsets as indicated in pLNs and mLNs. Numbers adjacent to outlined areas represent the percentage of cells in the area. Each symbol represents an individual throughout. Data are representative of four independent experiments. Data were represented as means ± SEM and analyzed with two-tailed unpaired *t* test. pLNs: peripheral lymph node. mLNs: mesenteric lymph node.

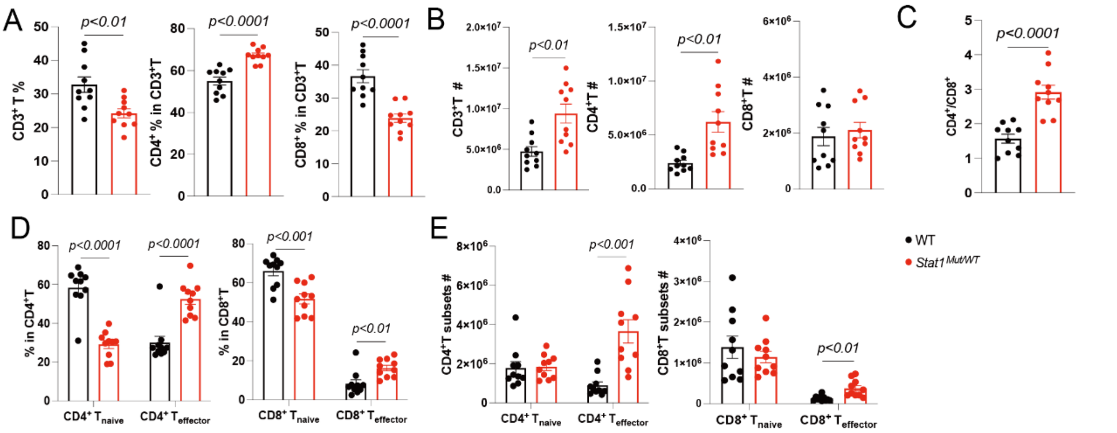


**Fig. S5 Expansion of CD4$^+$T cells in STAT1 GOF mice at age of 20-22 weeks.** (**A**) Percentage of CD3$^+$ in live cells, CD4$^+$ in CD3$^+$T, CD8$^+$ in CD3$^+$T cells in spleen. (**B**) Numbers of splenic CD3$^+$, CD4$^+$ and CD8$^+$ T cells. (**C**) Ratio of CD4$^+$ to CD8$^+$ T percentage. (**D-E**) Percentage (D) and number (E) of CD4$^+$ and CD8$^+$T subsets. Each symbol represents an individual throughout. Data are representative of four independent experiments. Data were represented as means ± SEM and analyzed with two-tailed unpaired *t* test.

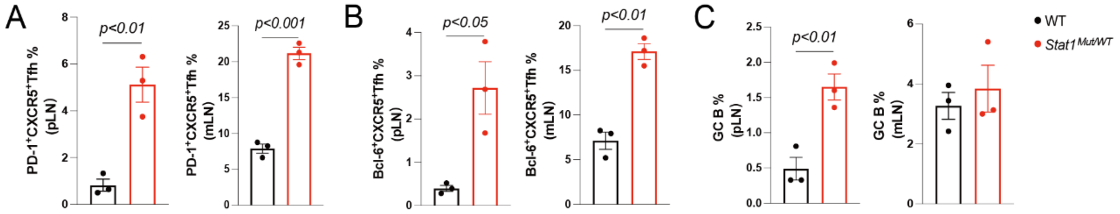


**Fig. S6 Expanded Tfh cells in lymph nodes of STAT1 GOF mice.** (**A**) Percentage of PD-1$^+$CXCR5$^+$Tfh (gated in CD4$^+$CD44$^+$Foxp3$^-$) in pLNs and mLNs. (**B**) Percentage of Bcl-6$^+$ CXCR5$^+$ Tfh (gated in CD4$^+$CD44$^+$Foxp3$^-$) in pLNs and mLNs. (**C**) Percentage of GC B cells in pLN and mLN. Each symbol represents an individual throughout. Data are representative of three independent experiments. Data were represented as means ± SEM and analyzed with two-tailed unpaired *t* test.

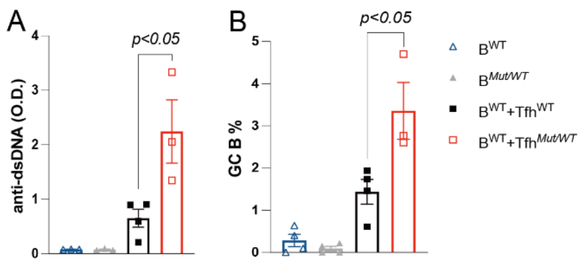


**Fig. S7 Tfh from STAT1 GOF mice induced autoantibody production.** (**A-B**) Bar graph showing O.D. value of serum anti-dsDNA antibody (A) and percentage of GC B cells gated in $B220^+$ cells (B). Each symbol represents an individual throughout. Data are representative of two independent experiments. Data were represented as means ± SEM and analyzed with two-tailed unpaired *t* test.

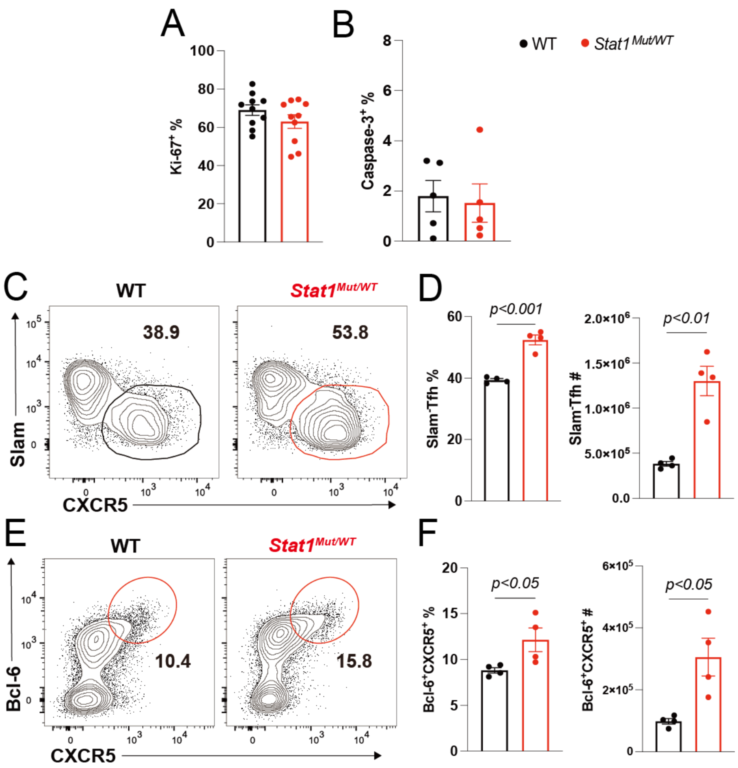


**Fig. S8 Expanded Tfh cells in STAT1 GOF mice infected with LCMV *Armstrong*.** (**A**) Percentage of Ki-67$^+$Tfh cells of mice at age of 20-22 weeks. (**B**) Percentage of Caspase3$^+$Tfh cells of mice at age of 20-22 weeks. (**C**) Flow cytometry plot of Slam$^-$CXCR5$^+$Tfh cells (gated in CD4$^+$CD25$^-$GITR$^-$CD44$^+$T cells) of mice infected with LCMV Armstrong. (**D**) Percentage (left) and number (right) of Slam$^-$Tfh cells of mice infected with LCMV Armstrong. (**E**) Flow cytometry plot of Bcl-6$^+$CXCR5$^+$Tfh cells of mice infected with LCMV Armstrong. (**F**) Percentage (left) and number (right) of Bcl-6$^+$CXCR5$^+$Tfh cells of mice infected with LCMV Armstrong. Numbers adjacent to outlined areas represent the percentage of cells in the area. Each symbol represents an individual throughout. Data are representative of three (A-B) or two (C-F) independent experiments. Data were represented as means ± SE and analyzed with two-tailed unpaired *t* test.

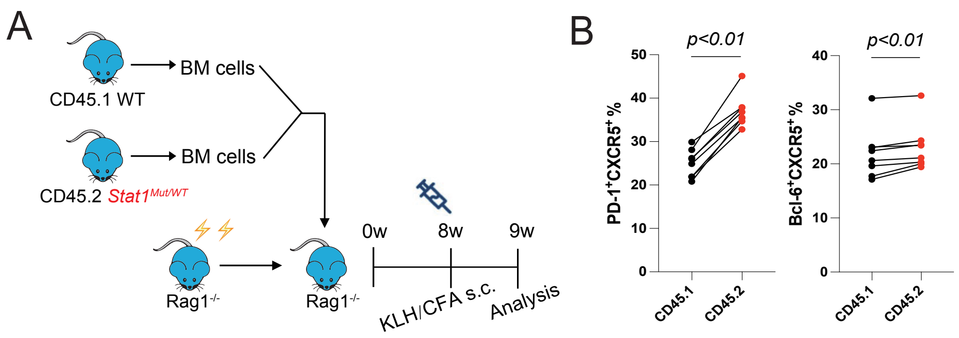


**Fig. S9 Intrinsic expansion of Tfh cells from STAT1 GOF mice.** (**A**) Schematic diagram of bone marrow chimera model. (**B**) Percentage of $CXCR5^{+}PD\text{-}1^{+}$ Tfh and $CXCR5^{+}Bcl\text{-}6^{+}$ Tfh from different donors. Each symbol represents an individual throughout. Data are representative of two independent experiments. Data were represented as means ± SEM and analyzed with two-tailed paired *t* test.

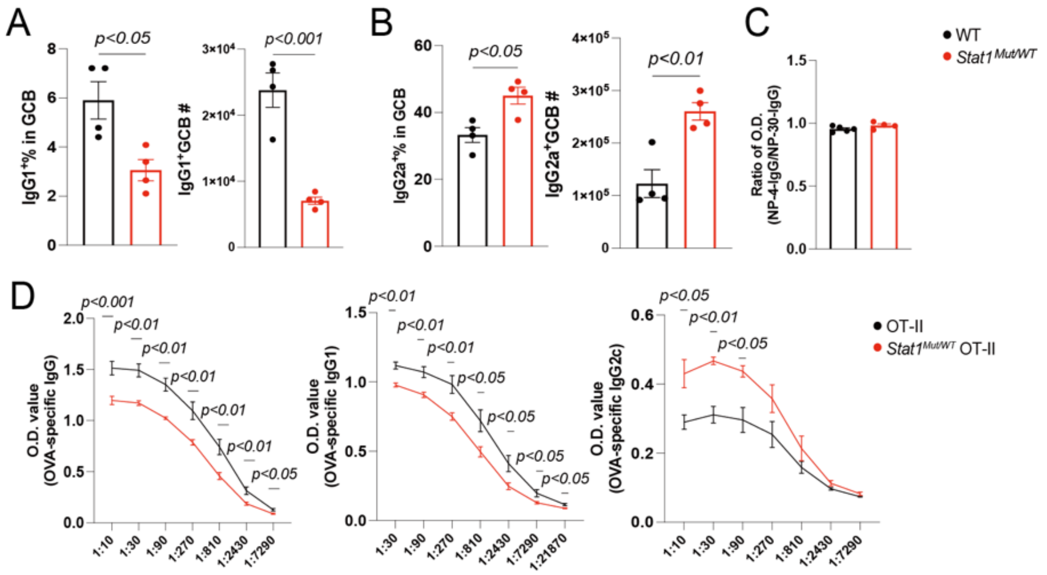


**Fig. S10 Defective humoral responses in different immune challenge, related to Figure 5.** (**A**) Percentage and number of IgG1$^{+}$GCB cells of mice following LCMV Armstrong infection. (**B**) Percentage and number of IgG2a$^{+}$GCB cells following LCMV Armstrong infection. (**C**) Ratio of NP-4-specific IgG to NP-30-specific IgG following NP-KLH/CFA immunization. (**D**) Total IgG and IgG subtype levels in serum of adoptive transfer model related to Figure 5. Each symbol represents an individual throughout. Data are representative of two (A-B) or three (C-D) independent experiments. Data were represented as means ± SE and analyzed with two-tailed paired *t* test.

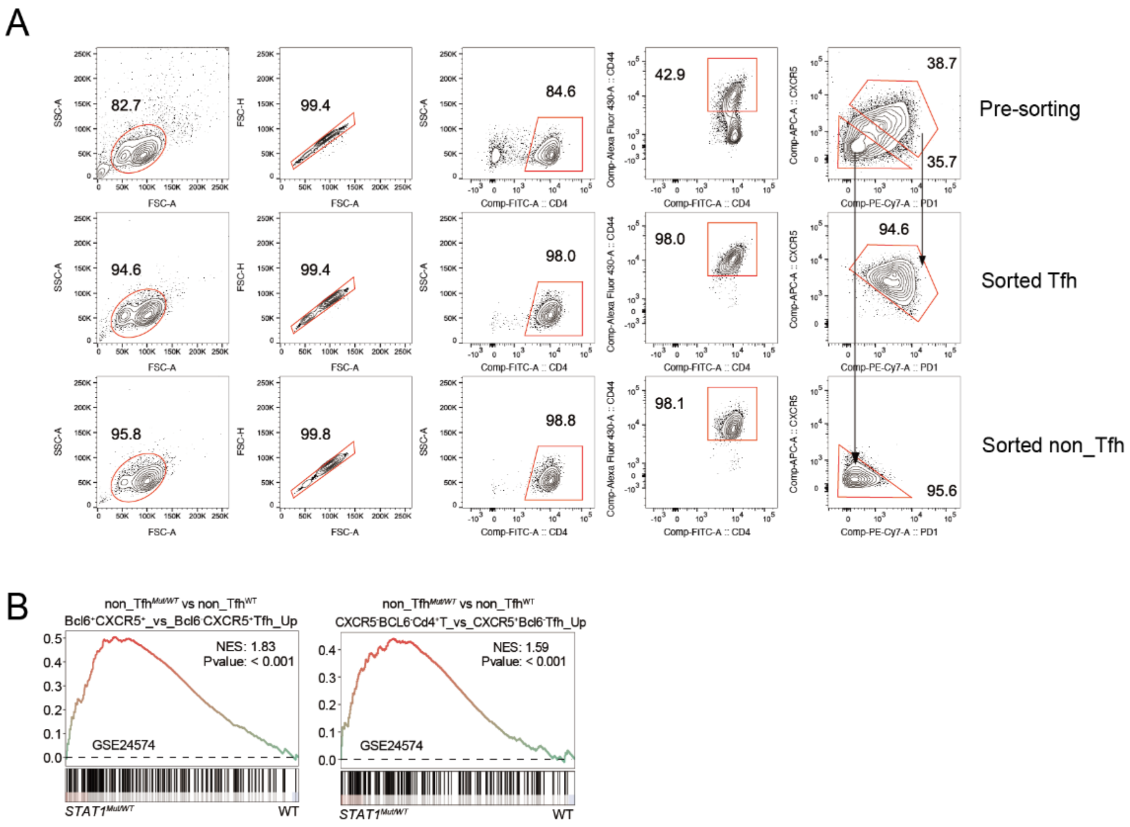


**Fig. S11 Cell sorting strategy and GSEA analysis, related to Figure 6.** (**A**) Gating strategy of Tfh and non_Tfh sorting using flow cytometry. Pre-sorting (upper), sorted Tfh (middle) and sorted non_Tfh (bottom). (**B**) GSEA plot for the indicated gene set showing NES and nominal *P* value.

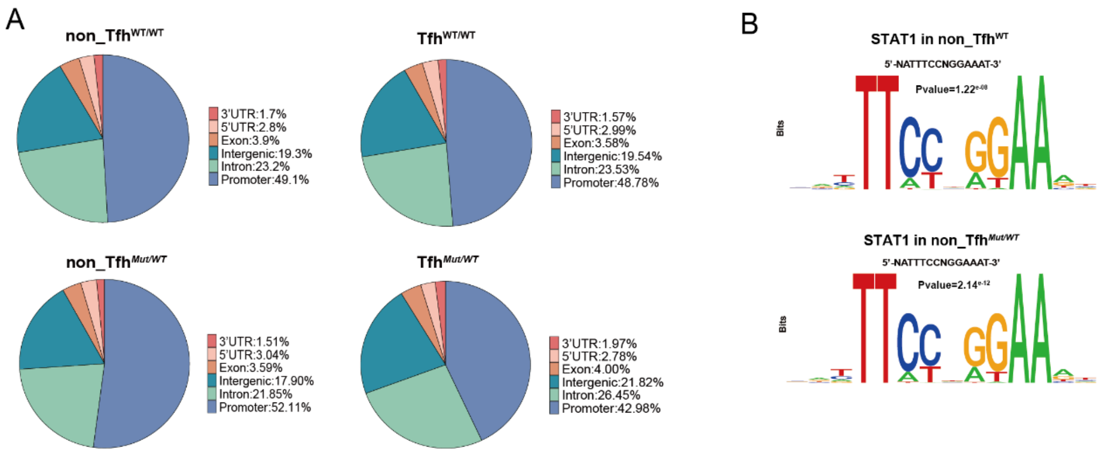


**Fig. S12 Distribution of STAT1 binding profile in gene loci**. (**A**) Pie graph showing the distribution of CUT & Tag peaks in different cells as indicated. (**B**) STAT1-binding motif in non_Tfh cells from indicated mice.

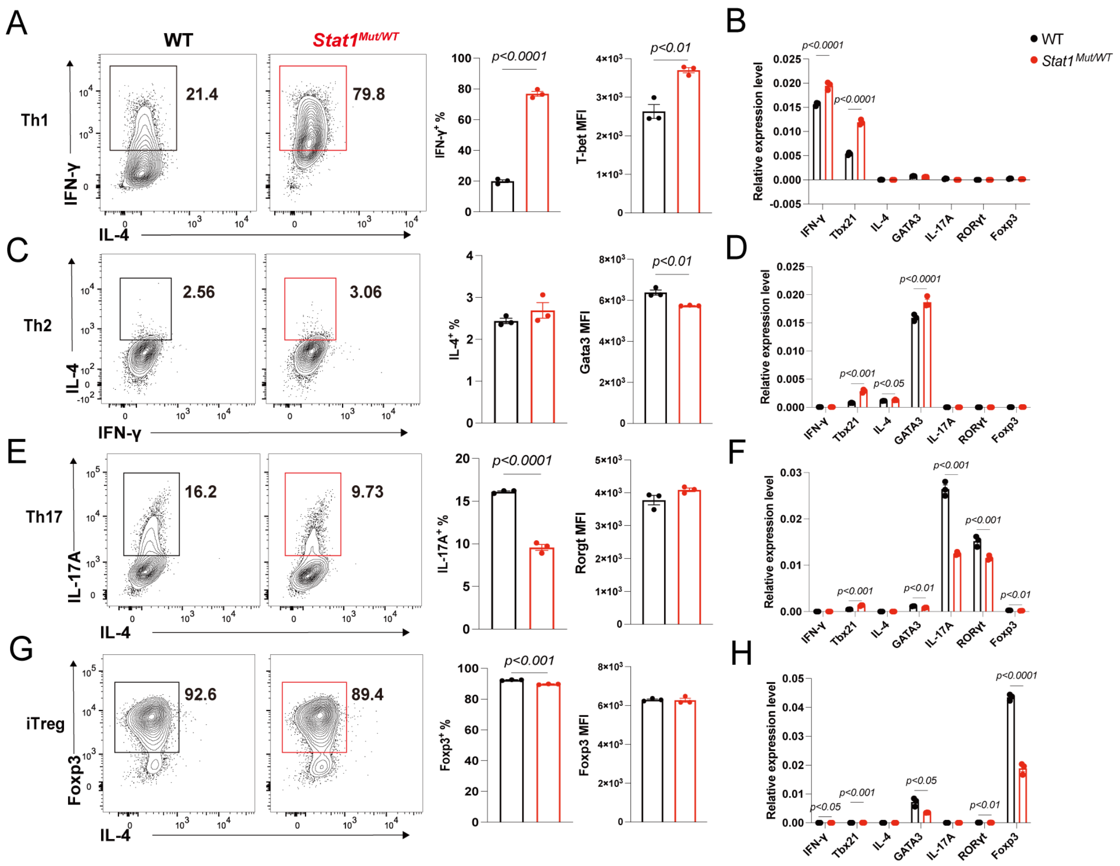


**Fig. S13 Dysregulated CD4$^+$T cells differentiation *in vitro*.** Isolated naive CD4$^+$T cells from WT and mutant mice were polarized under Th1(A-B), Th2(C-D), Th17 (E-F) and inducible Treg (iTreg, G-H) conditions for 3 days. Expression of signature genes were determined by intracellular staining and real-time RT–PCR. Each symbol represents an individual throughout. Data are representative of two independent experiments. Data were represented as means ± SEM and analyzed with two-tailed paired *t* test.

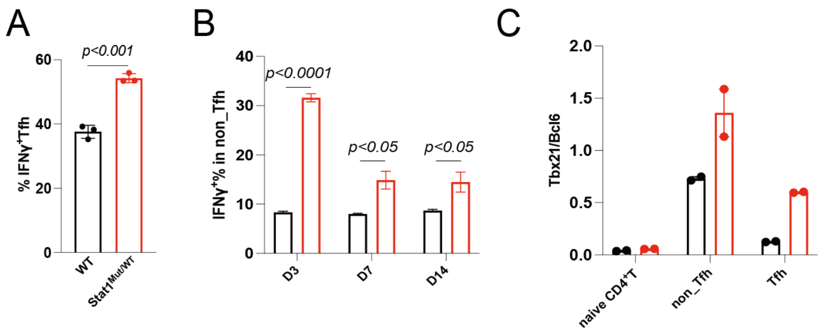


**Fig. S14 Tfh1 skewing phenotype in STAT1 GOF mice, related to Figure 8.** (**A**) Percentage of IFN-$\gamma^+$ in Tfh of mice at age of 6-8 weeks. (**B**) Percentages of IFN-$\gamma^+$ in non_Tfh of mice after immunization with KLH/CFA for indicated time. (**C**) Ratio of *Tbx21* to *Bcl-6* expression in indicated cells. Each symbol represents an individual throughout. Data are representative of two independent experiments. Data were represented as means ± SEM and analyzed with two-tailed paired *t* test.

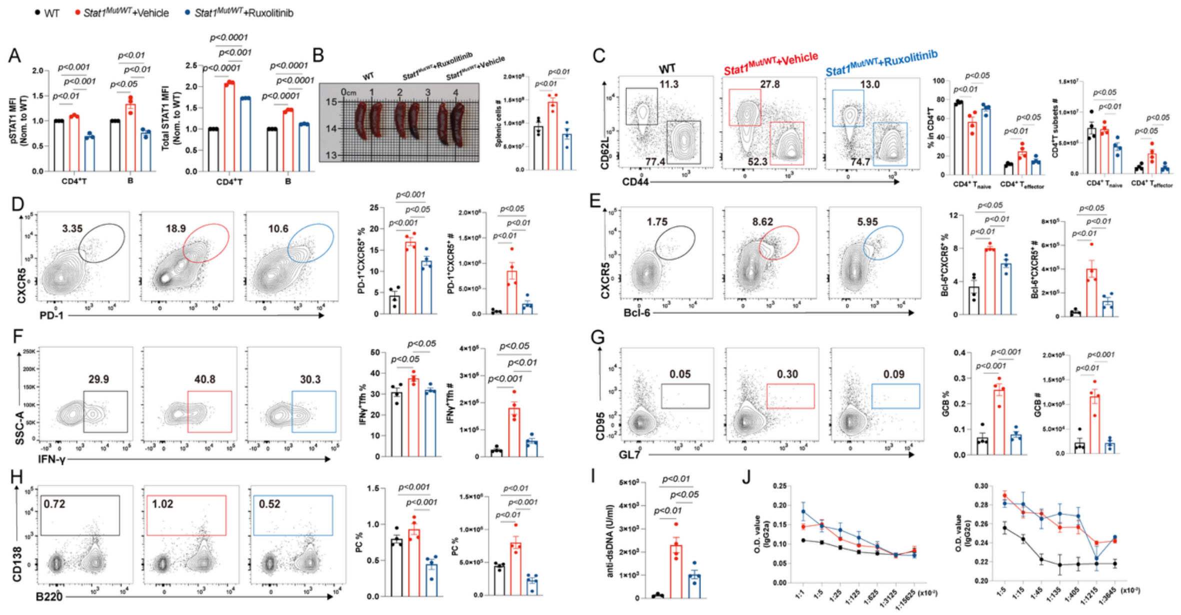


**Fig. S15 Autoimmunity phenotype in STAT1 GOF mice reverted by Ruxolitinib treatment**

8-week-old *STAT1*$^{mut/WT}$ mice were treated with ruxolitinib (50mg/kg per day) by oral gavage once daily for 2 weeks or received vehicle alone. After 2 weeks of administration, (**A**) the levels of phosphorylated STAT1 and total STAT1 in $CD4^+$T cells and B cells of the mice were determined. **(B)** Image of spleen (left) and number of splenocytes in different mice as indicated. (**C**) The percentage and number of $CD4^+$T subsets in different group. (**D-E**) Flow cytometry plot, percentage and number of $CXCR5^+PD\text{-}1^+$ T and $CXCR5^+Bcl\text{-}6^+$ T cells. (**F**) Flow cytometry plot, percentage and number of IFN-$\gamma^+$Tfh cells in indicated group. (**G-H**) Flow cytometry plot, percentage and number of GC B cells and $CD138^+$Plasma cells. (**I**) Serum anti-dsDNA antibody detected by ELISA. (**J**) IgG2a and IgG2c levels in serum of mice as indicated by colors. Numbers adjacent to outlined areas represent the percentage of cells in the area. Each symbol represents an individual throughout (n=4-5 mice per group). Data were represented as means ± SEM and analyzed with two-tailed unpaired *t* test.

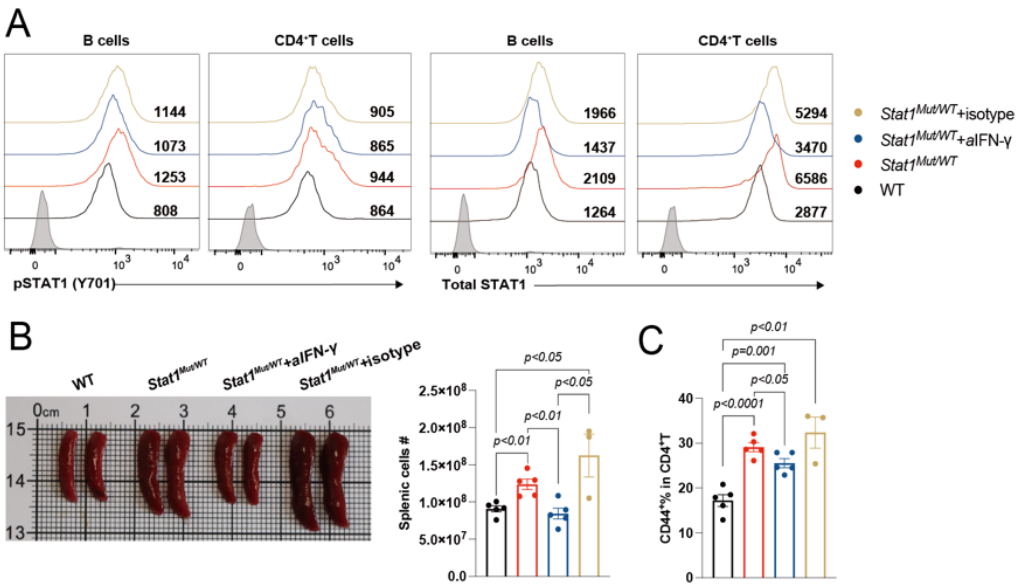


**Fig. S16 Alleviated STAT1 hyperactivity and CD4$^{+}$T cells activation following αIFN-γ treatment, related to Figure 8**.

(**A**) Histogram plot showing pSTAT1 (Y701) and total STAT1 in B cells and CD4$^{+}$T cells. (**B**) Image of spleen and number of splenocytes in different group as indicated. (**C**) Bar plot showing percentage of CD44$^{+}$ in CD4$^{+}$T cells. Each symbol represents an individual throughout (n=3-5 mice per group). Data were represented as means ± SEM and analyzed with two-tailed unpaired *t* test.
