## Supplementary figures and images for "Hyperactive STAT1 Promotes T Follicular Helper Type 1 Cell Differentiation to Trigger Autoimmunity"

### Graphical abstract.pdf

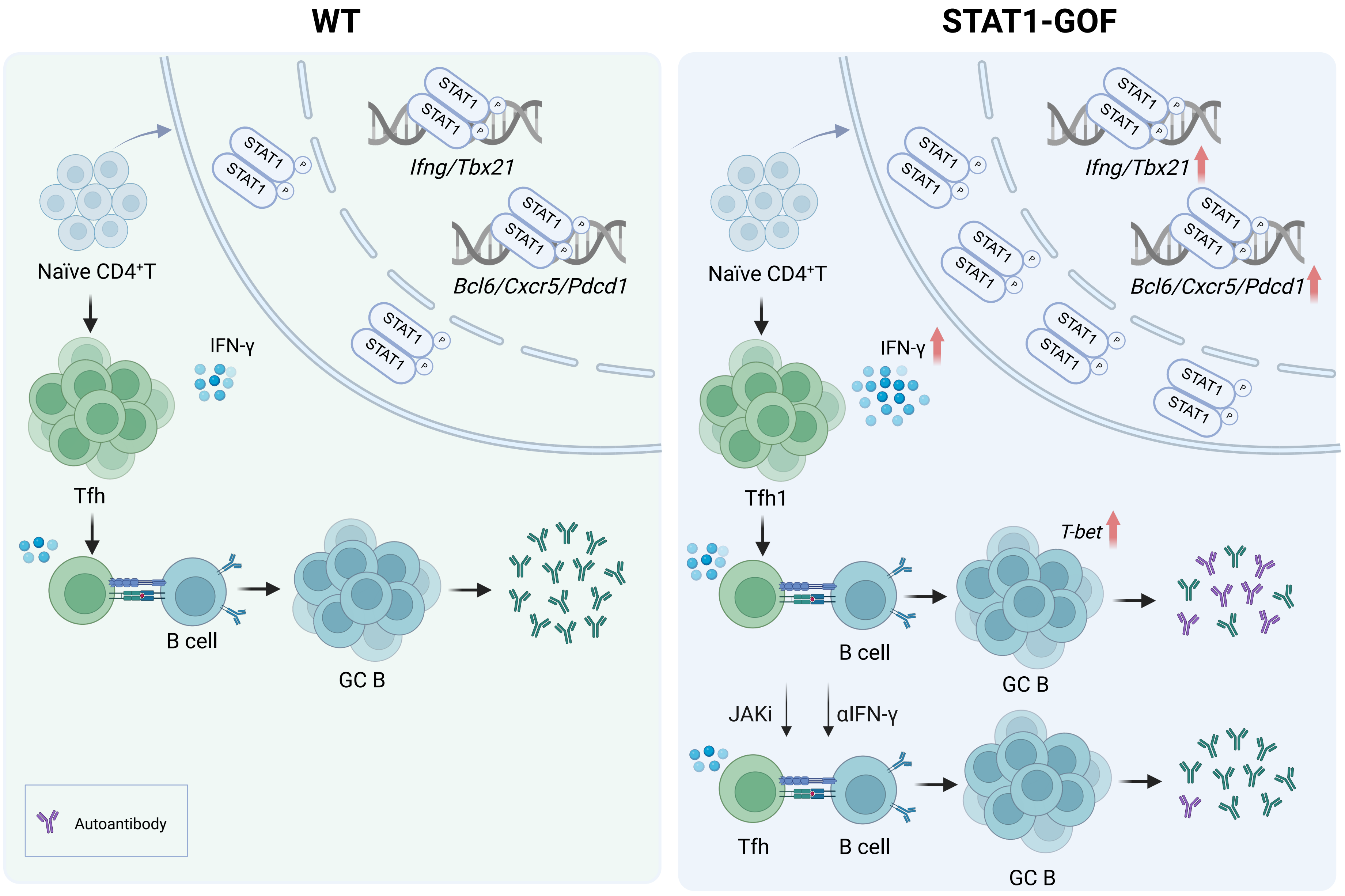
